# Artiphysiology reveals visual preferences underlying V4-like blur selectivity in a deep convolutional neural network

**DOI:** 10.1101/2019.12.24.886002

**Authors:** Saba Entezari, Wyeth Bair

## Abstract

Blurry visual scenes arise from many causes and image blur is known to be important for scene interpretation, yet there have been very few studies of the visual encoding of blur in biological visual systems or in artificial visual systems trained for object recognition. Recently, a study of single neurons in the visual cortex of macaque monkeys found that a significant fraction of neurons were more responsive to blurred visual stimuli than to sharply defined stimuli. This raises two questions: (1) what types of visual features in natural scenes might underlie blur selectivity in macaque cortex, and (2) can blur selectivity be found in artificial neural networks using the simple artificial stimuli previously applied in vivo? To answer these questions, we presented simple shape stimuli to the widely studied deep convolutional neural network (CNN) known as AlexNet and used deconv-net visualization to identify features that are critical for driving blur selective units. We found that a substantial number of units in the CNN were selective for blur and that several categories of blur selectivity emerged in early-to-middle processing stages. Prominent among these is a set of units selective for spatial boundaries defined by blur contrast. These blur-contrast boundary units may serve the important task of segmentation in natural photographic images and may relate to a large body of literature on second-order boundary detection. Our results lead to the prediction that such units could exist in the visual cortex but have yet to be well characterized and localized, and they provide direction for future neurophysiological tests of blur selectivity in vivo.

## INTRODUCTION

Blur is ubiquitous in natural images, arising from out of focus objects, cast shadows, shading on smooth surfaces, diffuse scattering of light through air and water and relative motion between the imaging system and environment. Thus, much information about a scene is carried by blur and low spatial frequency gradients (Elder, 2010), and it has been shown, for example, that observers can use blur to recover depth (Mather, 1997; Held et al., 2012). Blur is also important in artificial vision systems where spatial maps of blur are sought to aid scene interpretation (e.g., Ma et al., 2018). In image datasets used in classification contests, many categories contain photographs of objects in sharp focus, for example birds and insects, against blurry backgrounds, thereby making blur a key indicator of the location of the object to be classified. In spite of its importance, blur has received little direct attention in visual neurophysiology.

One recent study in the mid-level cortical area V4, part of the ventral, form-processing stream, found that some neurons believed to be tuned for the shape of object boundaries were most selective for intermediate levels of blur (Oleskiw et al., 2018). This study was carried out with simple shape stimuli that were blurred to varying degrees by Gaussian convolution, and thus it remains unknown what the selectivity of such blur-preferring neurons might be if tested with a broad range of natural images. To answer this question in vivo remains nearly impossible because of the severely limited number of visual stimuli that can be presented while single neurons are recorded in an awake primate. However, artificial neural networks are now available that offer sophisticated visual systems that have single-unit tuning similar to that of neurons in area V4 in terms of translation-invariant shape selectivity (Pospisil et al., 2018).

We therefore took the set of simple blurred stimuli used in the neurophysiological study of Oleskiw et al. (2018) and presented them to the same CNN tested by Pospisil et al. (2018) with the goal of determining (1) to what degree simple artificial stimuli are useful for detecting blur selective units in a deep network, and (2) what types of selectivity such units have when tested on natural scenes. We found that many units were identified as blur selective across all levels of the network and we defined a class of units that we refer to as blur-contrast boundary (BCB) units that respond best to oriented or shaped boundaries between regions of high and low blur levels. Our results raise the testable prediction that similar units may be prevalent in vivo, related to second-order boundary processing (Mareschal and Baker, 1998; Dakin and Mareschal, 2000; Baker and Mareschal, 2001; DiMattina and Baker, 2019), and they provide insight into improving simple stimulus sets for testing for blur selectivity.

## METHODS

### Visual stimuli

Our stimulus set was like that of Oleskiw et al. (2018), which was based on the design of Pasupathy and Connor (2001). It was generated from 51 simple shapes (Figure 1A) drawn at up to eight rotations each (Figure 1B; 45° increments), where rotations producing a replica of a previous rotation were omitted. Each of the resulting 362 rotated shapes was presented at seven levels of blur (Figure 1C), resulting in 2,534 unique stimuli. Blur level was quantified as the SD of the Gaussian used to convolve (blur) the unblurred (SD = 0) shape. We used blur levels of 0, 1, 2, 4, 8, 16 and 32 pixels (red labels in Figure 1C). Shapes were generated as bright achromatic forms (pixel value 128 for R, G and B) on a darker background (pixel value 0) and then blurred using an area-preserving convolution. The size of the shapes was quantified as the diameter of the large circle (second shape from left, top row, Figure 1A) before blurring, with all other shapes scaled to maintain the proportionality shown. We tested shape sizes of 31 and 155 pixels (the input image for the CNN was 227 by 227 pix) to accommodate units with a variety of receptive field (RF) sizes. Stimuli were presented in the center of the visual field of the network.

**Figure 1.**
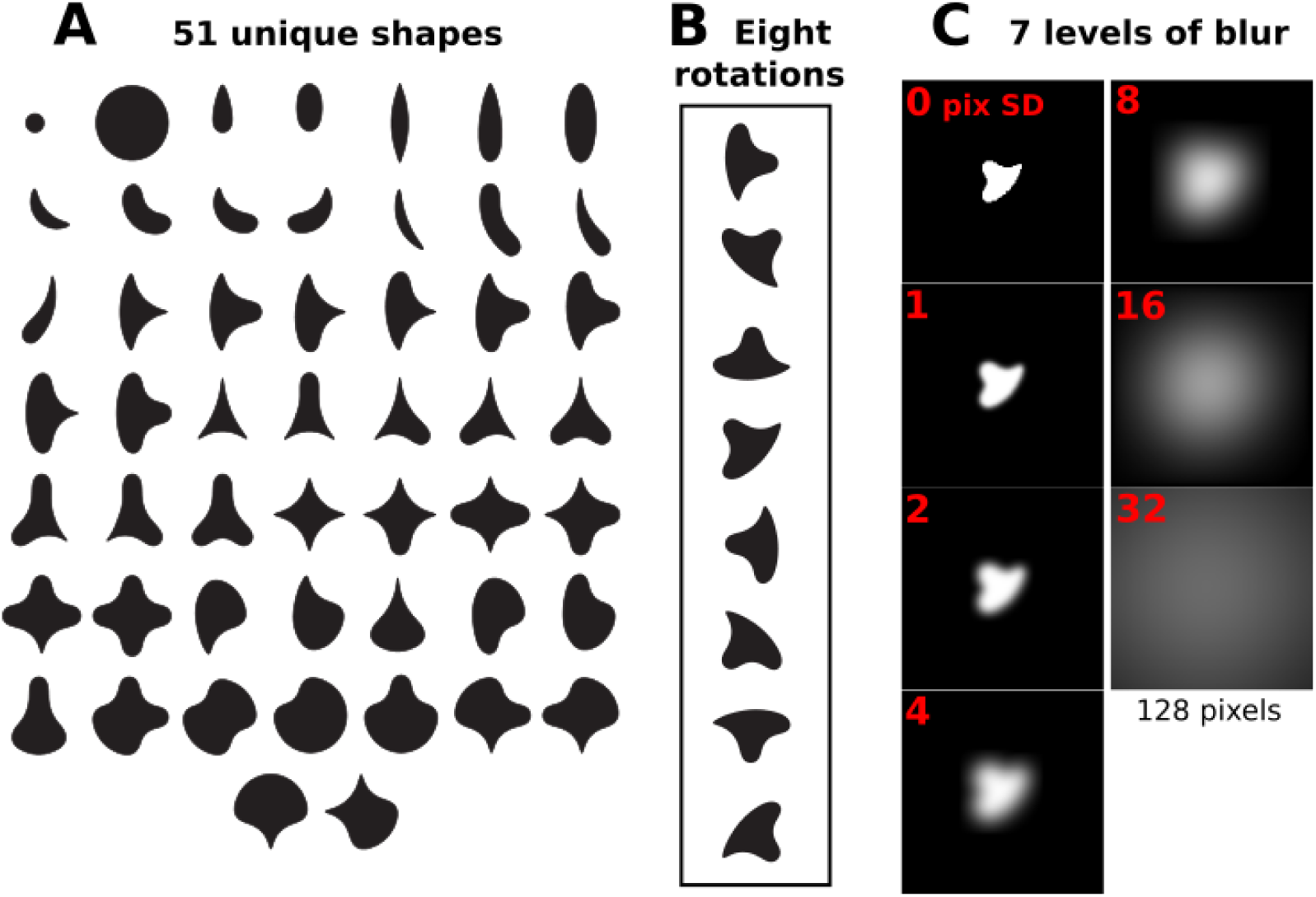
Visual stimuli. **(A)** The 51 unique shapes from Pasupathy and Connor (2001). **(B)** Each shape was shown at eight rotations (illustrated here for one example shape) or fewer to avoid repeating identical stimulus images. For example, the small and large circle stimuli (upper left) were shown at only one rotation, because all rotations are identical. **(C)** Each shape was presented at 7 levels of blur. Shapes were drawn as white on a darker (mid gray) background. The background is rendered here as black to enhance contrast (see Methods) and only the central 128 pix square region is shown (the full stimulus image is 227 pix square).

### Criteria for identifying potentially blur selective units

Our fundamental metric is b_max_, defined as the blur factor associated with the maximum response across all shapes and blur levels. Based on b_max_, we defined two criteria for considering a unit to be potentially blur selective. Our “lax” criterion was that 0 < b_max_ < 32, meaning that bmax must be an intermediate (not extremal) blur factor, and that the mean response of the unit across the 362 shapes at b_max_ was larger than its mean response for no blur (b=0) and larger than its mean response at the highest blur factor (b=32). Our “strict” criterion maintained the condition that b_max_ was intermediate and further required that the mean at b_max_ exceeded even the maximum responses at b=0 and b=32. We tested other metrics for blur selectivity based on response dynamic range, but found results similar to those reported here.

### The convolutional neural network (CNN)

We analyzed a pre-trained version of AlexNet available on the Caffe machine learning website (Jia et al., 2014). This is the same network that was shown by Pospisil et al. (2018) to contain units that have V4-like shape selectivity and translation invariance. We examined the description of the architecture, downloaded the weights and biases, and reproduced the network in custom C code. We used test images to establish that responses from our C implementation were essentially identical to those from the Caffe software. The main features of the network are described in the original publication (Krizhevsky et al., 2012) and details of our implementation are described in Pospisil et al. (2018, see Figure 2 therein for architecture and unit RF sizes). Briefly, the network has eight major layers where the first five are convolutional layers (Conv1, Conv2, …, Conv5) and the last three layers (FC6, FC7 and FC8) are “fully connected” meaning that each unit receives weighted signals from all units in the previous layer and all units are unique. We tested only the unique units in the network which have counts as follows: 256 in Conv2 and Conv5, 384 in Conv3 and Conv4, 4096 in FC6 and FC7 and 1000 in FC8. The final FC8 layer units have a 1-to-1 correspondence to 1000 image categories. We did not test Conv1 because it is trivial. Here we focus on the signed outputs of the first sublayer within each major layer. We also analyzed signals following rectification, max pooling and normalization sublayers and found largely similar results.

**Figure 2.**
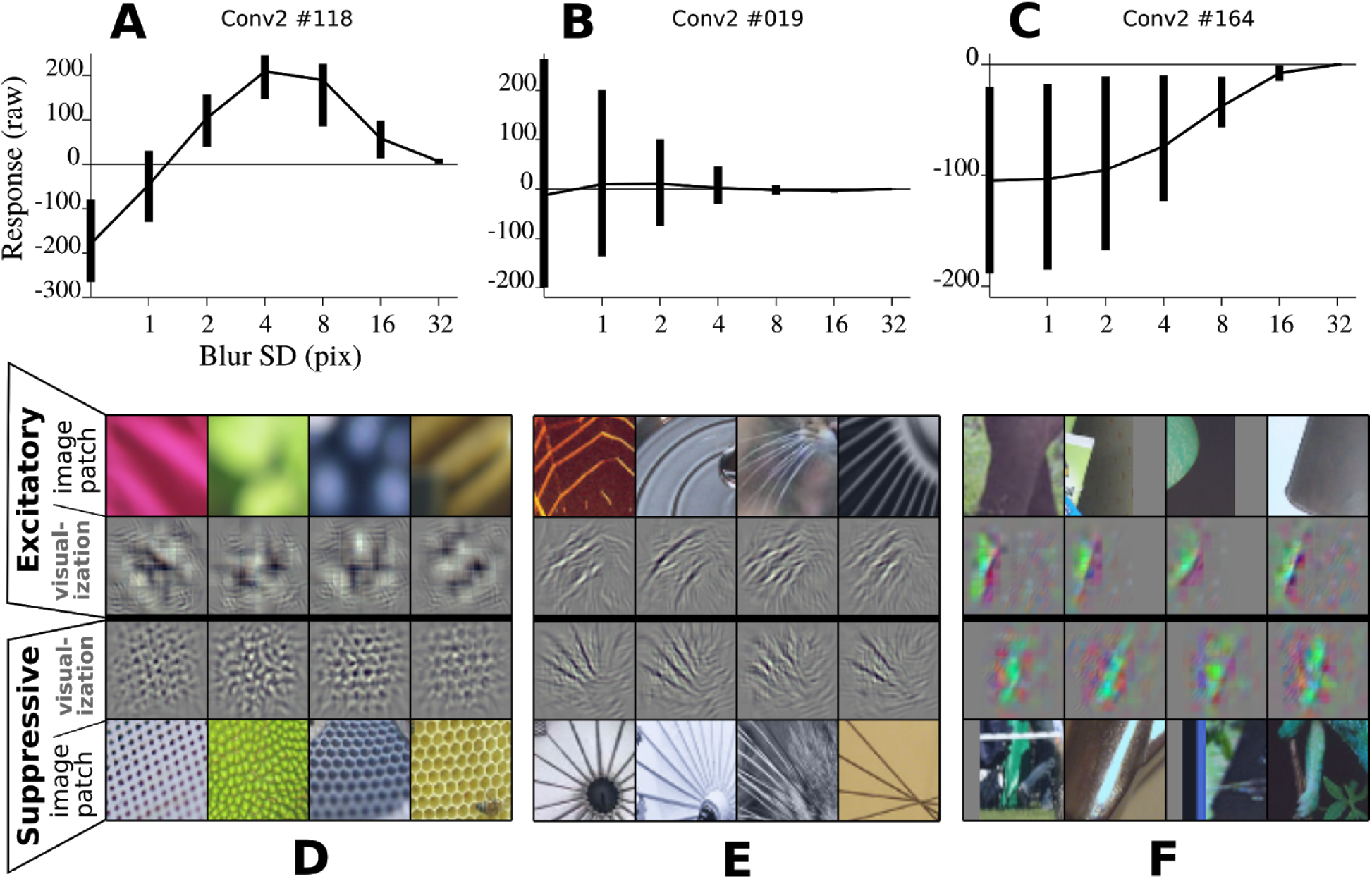
Response range as a function of blur. Three units in layer Conv2 exemplify different patterns of response as a function of blur. The thick vertical bars in the upper graphs indicated the response range from the minimum to maximum value across 362 shapes at each blur level. The thinner lines connect the mean response values at each blur level. **(A)** Unit #118 responds most strongly to shapes with intermediate blur (b=4), and is strongly suppressed by unblurred (b=0) shapes. **(B)** Unit #119 is highly selective for unblurred shapes, being most strongly excited (positive activations) by some shapes but most strongly inhibited (negative activiations) by others. **(C)** Unit #164 is suppressed strongly and selectively by unblurred shapes, and this selective suppression becomes progressively weaker (responses compress upwards toward zero) at middle to high blur levels. **(D)** For the example unit in (A), the most excitatory four image patches are shown in the top row. In the second row, the visualization of critical features in each image patch is shown. At the bottom, the four most suppressive image patches are shown for the same unit. Directly above the patches, in the 3rd row, the visualization of critical features is shown. **(E)** Same format as (D), images and visualization for the unit in (B). **(F)** Images and visualization for the unit in (C). In the second and third excitatory image patches and the first and third suppressive image patches, gray boxes have been added where the potential RF extends beyond the input image.

The version of AlexNet used here is similar to the original in that the Conv layer units were split in half in some layers to facilitate the training process. This resulted in one set (the first half of the kernels) developing largely achromatic sensitivity in Conv1 (see Figure 3 in Krizhevsky et al., 2012) and the second half of the Conv1 units being predominantly chromatic. This tendency to differ with respect to chromatic sensitivity was propagated through the Conv layers. In one analysis, we analyze the units in these two sets separately.

**Figure 3.**
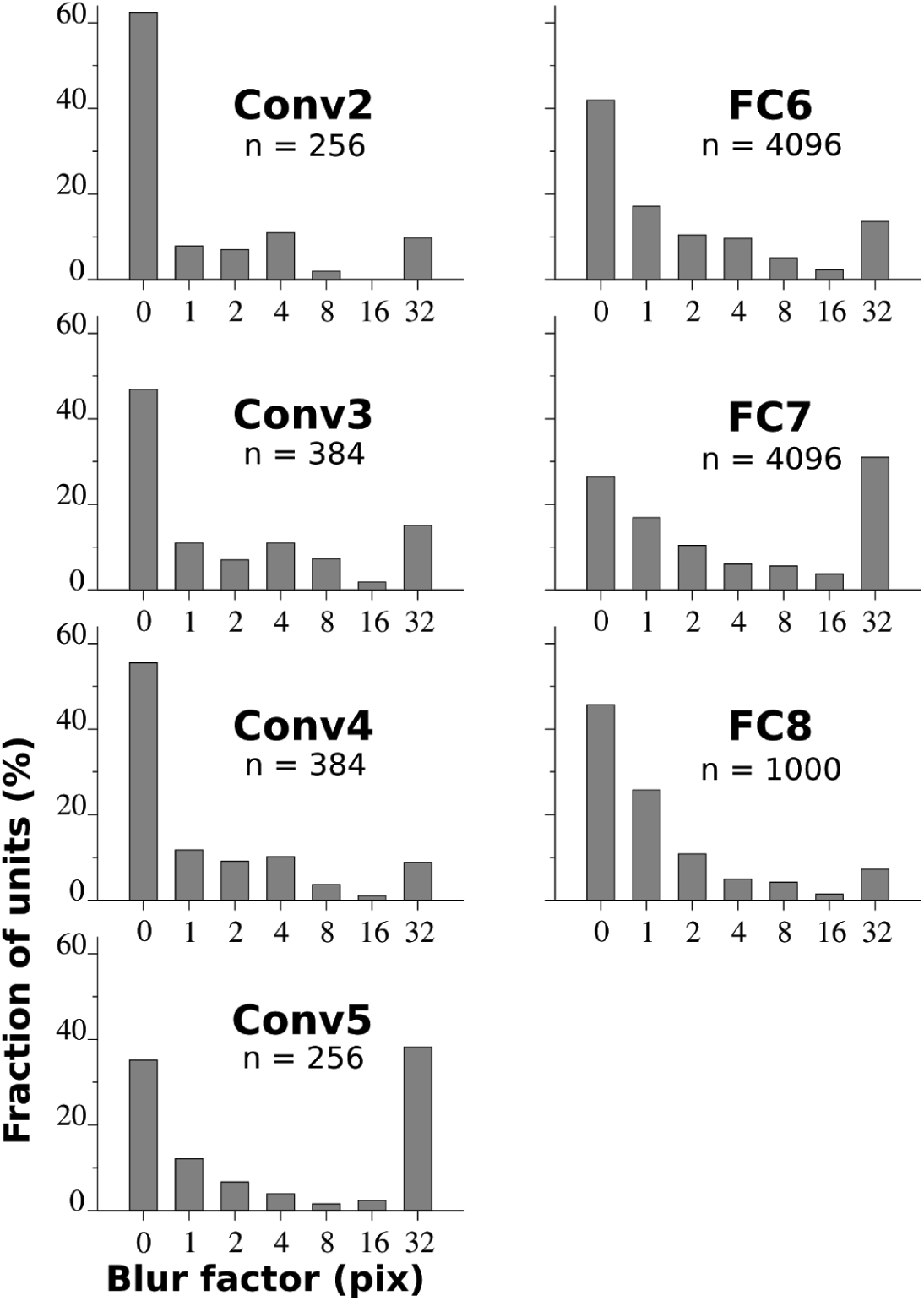
Distribution of blur factors, b_max_, associated with the maximal response of each unit. Each panel shows the bmax distribution for a given layer in the CNN. The number of unique units is indicated. The stimulus size was 31 pix and was presented as white on mid-gray.

### Visualization

To determine which features in natural images drove individual units in the CNN to their highest (most positive) and lowest (most negative) activation levels, we searched through a standard test set of 50,000 images for AlexNet. For convolutional layer units, we considered the responses at all x-y locations for each unique kernel. Layer Conv2 consisted of a 27 by 27 unit spatial grid, therefore across all images, responses to 36.45 million image patches were examined for this layer. We then took the 10 most excitatory (preferred) and ten most suppressive (anti-preferred) image patches (or entire images for the FC layers) and carried out a deconv-net visualization (Zeiler and Fergus, 2013). This method essentially distributes the responses of a unit to a given image back onto the set of weighted inputs to that unit dividing the response proportionally by the weights. Negative values are suppression, the normalization function is overlooked, and the grid-position within a max-pool unit that achieved the maximum on the feedforward pass was the only line on which the response was pushed back to the previous layer. This resulted in an RGB image that could be examined to provide a qualitative sense of what features within an image drove the unit to such a large positive or negative value. We refer to these images as visualizations, and present them directly next to the image patches from which they were derived.

## RESULTS

### Detecting blur selective units in a shallow layer

We presented a standard set of shape stimuli with different levels of blur to the units in AlexNet. To assess whether a unit was potentially blur selective, we examined how the range of responses across the set of 362 shape stimuli varied with blur, b. For a representative blur-selective unit (Conv2 #118), Figure 2A shows that responses were negative for all shapes at b=0 (thick vertical bars indicate full range of responses, thin line connects mean responses over all stimuli), but even a small amount of blurring (b=1) caused some stimuli to evoke positive responses. At intermediate levels of blur, responses were positive to all shapes and the maximum response occurred at b=4. The highest blur level (b=32) was associated with a very narrow range of response values just above zero, consistent with the notion that very blurry shapes no longer have distinctive features that excite or suppress unit activation. A second example unit (Figure 2B) was not selective for blur—its maximum response and maximum dynamic range (the difference between the maximum and minimum responses) occurred for b=0, and successively higher levels of blur simply narrowed the response range around zero. A third example unit (Figure 2C) was suppressed (had negative responses) to all of our shape stimuli at all blur levels except at the maximum blur level (b=32), where responses were narrowly distributed around zero. While the maximum response technically occurs for b=32, it would not be useful to categorize this unit as blur selective on this basis. This pattern of results is better interpreted as there being no stimuli in our set with features matching the selectivity of the unit (a color contrast boundary as demonstrated next).

After testing all units in layer Conv2 with artificial shape stimuli, we searched within a set of about 36.5 million image patches to identify the top 10 preferred (most excitatory) and bottom 10 anti-preferred (most suppressive) natural stimuli for each unit. Because Conv2 is an early layer with limited RF size (at most 51 x 51 pixels), simply looking at the preferred and anti-preferred patches can often reveal the selectivity of the unit, but we also applied deconv-net visualization (Zeiler and Fergus, 2013) to the chosen image patches to more precisely determine the features that drove very large positive and negative responses. This is demonstrated in Figure 2D for the example blur-selective unit. The top four excitatory image patches are shown in the top row, and the feature visualization is shown in the second row (for full images from which the patches arise, see Figure A1 for unit Conv2-#118 in the Appendix). This unit prefers blurry regions across its entire RF and is most suppressed by regions of regular, high spatial frequency (SF) texture (lower row shows anti-preferred image patches). The visualizations (middle rows, Figure 2D) highlight the achromatic texture within the image patches. For the second example unit, the visualization reveals that the preferred stimuli are thin lines at a particular orientation, and the suppressive stimuli are thin lines oriented orthogonal to preferred. This is consistent with the observation that unblurred shapes can drive high or low responses (Figure 2B, b=0), presumably via high frequency edges with particular alignment, whereas any blurring of the shapes reduces the power at high SF. For the third example unit, the visualization shows that the preferred stimulus is an oriented chromatic boundary between a bright, greenish and a dark, reddish region, whereas the anti-preferred stimulus is a spatially offset greenish stripe. This is consistent with the observation from Figure 2C, that essentially none of the shape stimuli evoke a positive response.

To estimate the frequency of potential blur selective units in Conv2, for each unit we computed a frequency histogram of b_max_, the blur factor corresponding to the maximum response across all shapes. The histogram for the 256 units in Conv2 (Figure 3, upper left) reveals that most units (62%) respond best for no blur (b=0; cf. Figure 2B) and 10% respond best at maximal blur (cf. Figure 2C). The remaining 28% had maximum responses at intermediate blur, suggesting that suppression from high SF channels in the previous layer (Conv1) is prevalent within the Conv2 layer. Histograms of b_max_ for the deeper layers (Figure 3, labels indicate layer) show that the overall trend is similar across all layers, with the notable exception that for Conv5 and FC7, more units have b_max_ = 32 than any other value. Further analyses (not shown) suggest that this is part of a trend in which suppression alternates, being stronger in odd layers (3, 5 and 7) and weaker in the even layers. We carried out visualization for units that had intermediate b_max_ values and found a variety of selectivity patterns that were associated with blur, but we also found many units for which the association to blur was not directly apparent. We therefore developed two criteria to try to identify blur selective units with greater certainty, as follows.

Based on unit responses to our shape set, we defined two criteria for blur selectivity that we refer to as “lax” and “strict”. The lax criterion requires the mean response at b_max_ to be greater than the means at b=0 and b=32. The strict criterion requires the mean at b_max_ to be greater than all responses at b=0 and b=32. We initially examined Conv2, where RFs are small, thus we chose a shape size of 31 pix (Figure 4A, black outlined shape in center) that allowed shapes to fit within the maximal possible RF for Conv2. To examine the influence of shape size, we also tested larger shapes that were larger but that fit within the maximal RFs for Conv5 and deeper. The fraction of units in each layer that satisfied the lax and strict criteria is plotted against layer for small stimuli in Figure 4B. In Conv2, 28 units (11%) satisfied the lax criterion and only seven of those satisfied the strict criterion. For those 28 units, the amount by which the mean at b_max_ exceeded that at b=0 and b=32 is plotted in Figure 5A, where red points indicate those that satisfied the strict criterion. This graphic reveals which units have the most prominent peaks. The right-most point corresponds to unit #118, our first example unit (Figure 2A), the second and third most rightward points are units #058 and #006, which are similarly blur selective based on visualization (see Appendix Figures A4 and A5). For these three units, all of the top ten image patches are out of focus regions in photographs where a spatially offset object is in sharp focus, with the exception that one photograph is entirely out of focus. Of the remaining four strict-criterion units, two are similar to those discussed, but two others are qualitatively different, as described next.

**Figure 4.**
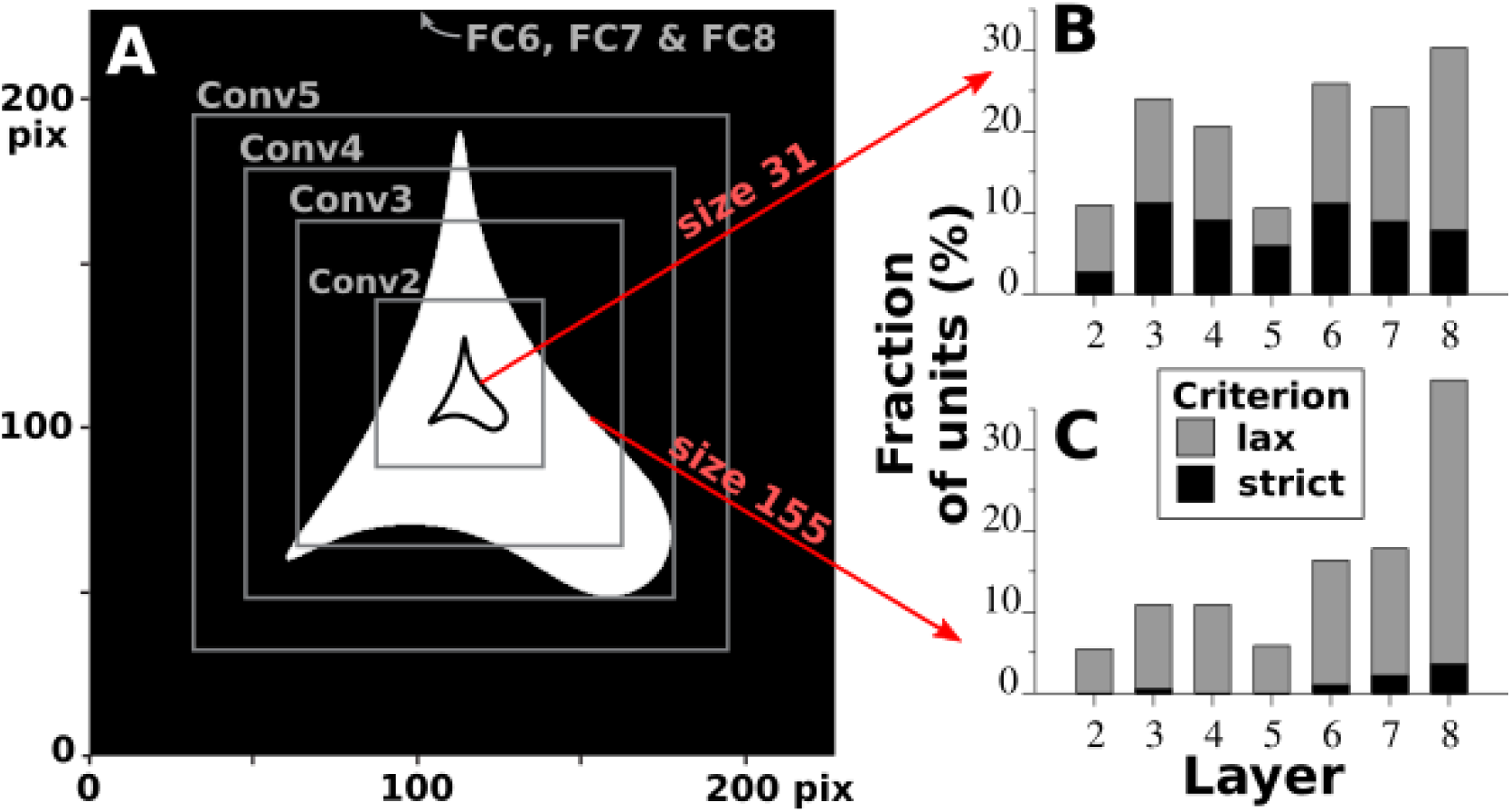
Fraction of units meeting two criteria for potential blur selectivity at two sizes. **(A)** Two stimulus sizes, 31 pix and 155 pix, are depicted (large white shape and small black outline, respectively) within the full input image space (black square region, 227 pix wide). The maximal possible receptive fields of units for layers from Conv2 to Conv5 are indicated by gray square lines. The maximal possible receptive field for FC6 to FC8 is the entire field. **(B)** For size 31 pix stimuli, the fraction of units that met the lax criterion (gray bars) and the subset of those that met the strict criterion (black bars). See Methods for criterion definitions. **(C)** For size 155 pix stimuli, the same as in (B).

**Figure 5.**
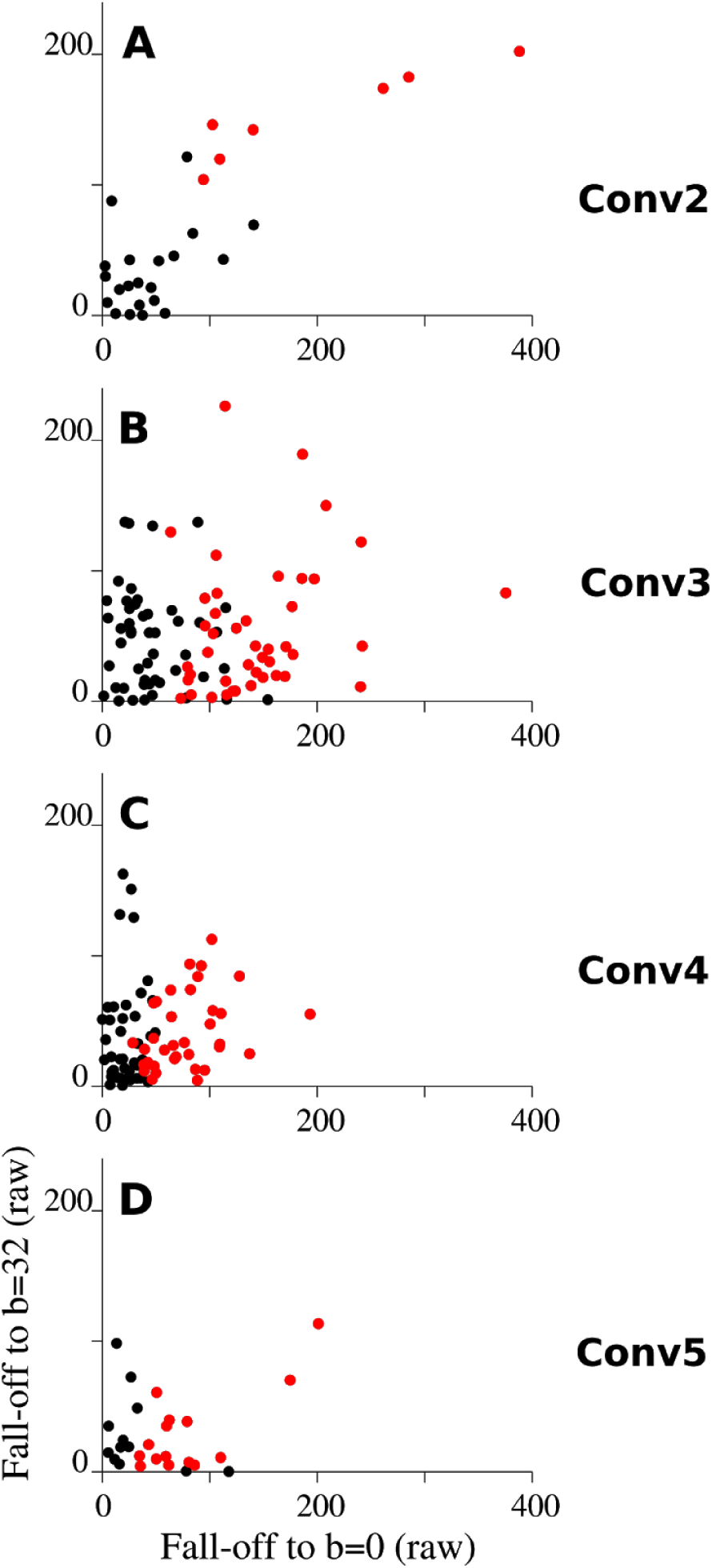
The amplitude of the peak of blur tuning curves for potentially blur selective units. For a subset of units in each Conv layer, a panel plots the mean at b_max_ minus the mean at b=32 (y-axis) against the mean at b_max_ minus the mean at b=0 (x-axis). Points are shown only for the subset of units for which both the x- and y-values were greater than zero (our lax criterion for potential blur selectivity). Points shown in red indicate units for which the mean at b_max_ was greater than all responses at b=0 and b=32 (our strict criterion).

The visualization for unit #077 (Appendix Figure A6) revealed a different selectivity pattern, where an oriented gradient of shading was often preferred, rather than simply an out of focus background. For example, gradients of reflected light on rounded surfaces appear on the vacuum and projector (Figure A6, 2nd and 4th top images), and low frequency gradients from light to dark (top to bottom) appear in the other images. This type of surface-generated or lighting-generated gradient selectivity occurs in other units and other layers.

The seventh unit to meet the strict criterion, #041 (Appendix Figure A7), revealed another type of tuning: selectivity for a boundary defined by blur contrast. We will refer to such units as blur-contrast boundary (BCB) units. All of the preferred image patches contain a blurry or blank region in the lower left against a high frequency textured region to the upper right. The consistency of the contrast between the textures is highlighted in the visualizations (2nd row, just below “Excitatory patches”, Figure A7). Importantly, the anti-preferred patches show the opposite pattern of selectivity—texture to the lower left and blurry or blank regions to the upper right. In this example unit, luminance may also play some role, as the preferred patches tend to have a light to dark gradient (lower-left to upper-right), which is reversed for the anti-preferred patches. Although this was the only BCB unit identified with our strict criterion, several others were among the 28 that met the lax criterion. For example, unit #027 (Figure A8) preferred a diagonal boundary between high SF texture at the lower left and blurry regions (particularly those consistent with rounded forms) to the upper right. Other units meeting the lax criterion are #103 (Figure A9), #099 and #080.

It is important to consider how these BCB units are detected by criteria based on having a stronger response for blurred than for non-blurred stimuli. If a unit prefers blur in one part of its RF and sharp focus in another, then it is possible that blurring a stimulus would increase the response in part of the RF while decreasing the response in the other part. Thus, in principle, BCB units should exist that are not detected by an overall peak at intermediate blur factors. We therefore carried out an exhaustive visualization of all 256 units in Conv2 to determine subjectively which ones appeared to be BCB based on spatial opponency and blur opponency. On this basis, we estimate that there are 36 units (14%) that are primarily BCB selective and another 16 units (6%) that are secondarily BCB selective. One such unit not detected by our criterion was #115 (Figure A10), which preferred sharply focused targets at the lower left against blurred backgrounds to the upper right. Another undetected unit was #203 (Figure A11), which preferred small red targets in the lower left against a featureless whitish field to the upper right. The response range plots for both of these units (Figure 6A and B) suggests that the units were generally suppressed by any amount of blur, yet this is in principle consistent for BCB units. This demonstrates why our simple shape set is insufficient to identify all units involved in processing blurry parts of scenes, and this may have important implications for in vivo studies as well (see Discussion).

**Figure 6.**
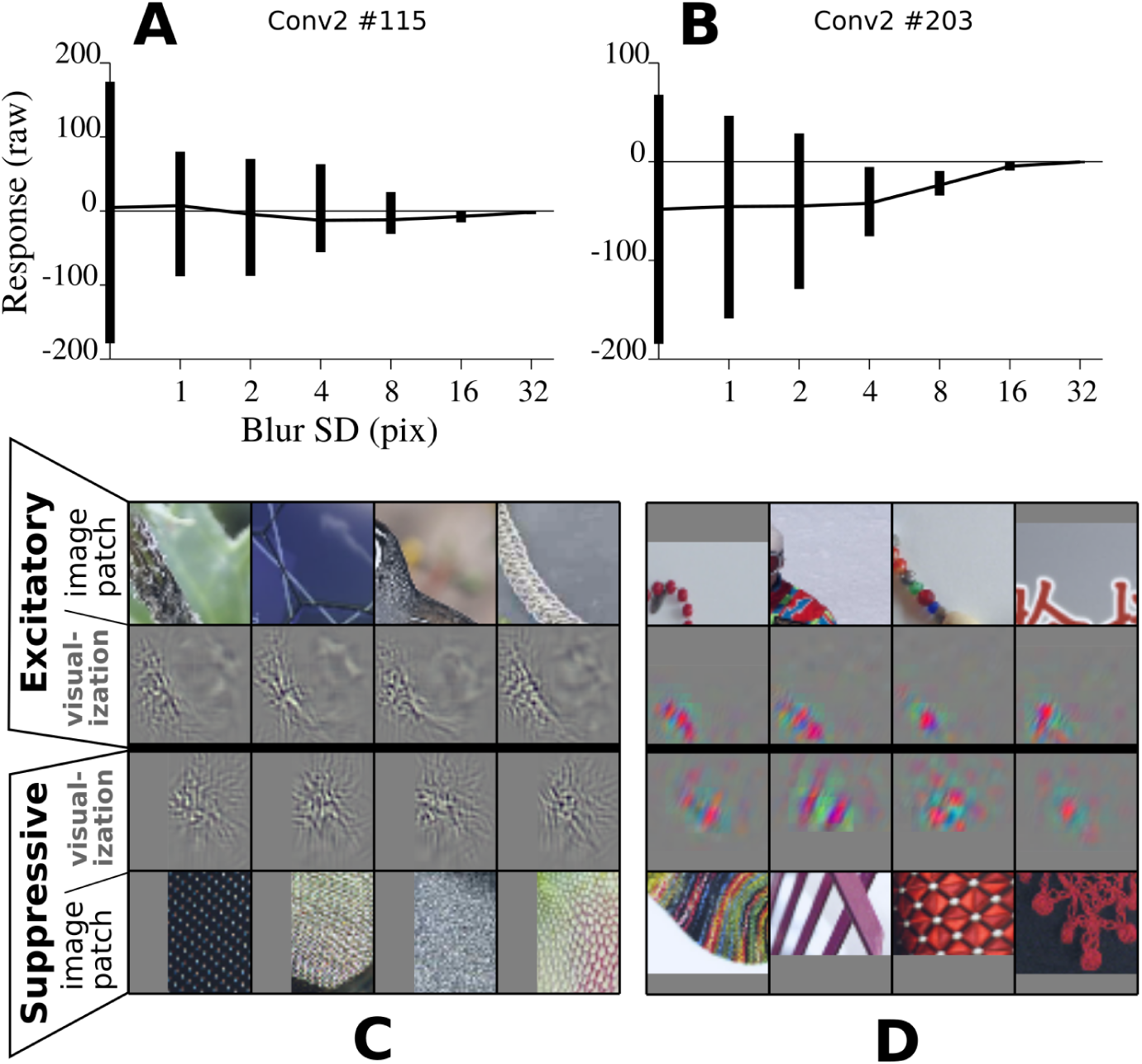
Response range plots for BCB units that were not detected by our blur criteria. **(A)** Response range plot for Conv2 #115. Same format as Figure 2A. **(B)** Response range plot for Conv2 #203. **(C)** and **(D)** The five most excitatory and inhibitory images and their deconv-net visualization for the units shown above in (A) and (B), respectively. Gray boxes are added where the potential RF extends beyond the input image. In the network, such regions are padded with all zeros and thus approximate featureless mid-gray regions.

Based on our examination of Conv2, it appears that a substantial fraction of units are involved in processing blur in images, including blurry out of focus regions, blur-defined boundaries, low frequency oriented gradients, broad rounded forms and transitions to blank regions. We next consider deeper layers.

### Blur selective units in deeper layers

The next two layers, Conv3 and Conv4, contained a substantially larger fraction of units that met our blur criteria. Whereas 2.7% of Conv2 units met our strict criterion for blur selectivity when tested with small shapes, 11% and 9% of units met this criterion in Conv3 and 4, respectively (Figure 4B). For nearly all such units, the visualization clearly showed blur-related features in the preferred images. Visualizations for the two rightmost points in the scatterplot for Conv3 (Figure 5B) reveal a unit, Conv3 #159, that prefers blurry or large rounded regions (Figure A11) and a unit, Conv3 #052, that prefers extended regions of fur (Figure A12). A substantial fraction of units meeting the strict criterion were BCB units, for example, Conv3 #048 (Figure A13). Unit Conv3 #302 preferred blank regions (Figure A14), highlighting the fact that blank regions and blur are difficult to distinguish. In Conv4, the right-most point in Figure 4C corresponds to unit #177 (Figure A15) that prefers relatively featureless or blurry regions with some chromatic selectivity contained within a boundary and that is suppressed by high SF and high contrast black-and-white circular patterns. Although the RFs are larger and the units are selective for more complex combinations of features, these units all build upon the types of blur selectivity observed in Conv2.

In Conv5, unit #026 (corresponding to the right-most point in Figure 5D) is selective for rounded forms (Figure A17), building on the gradient selectivity seen above. Unit #099 is selective for regions of background blur surrounded by sharp forms and is suppressed by sharply focused objects surrounded by blur (Figure A18), consistent with the integration of double-opponent (spatial and blur opponent) inputs from BCB units observed in shallower layers. Unit #039 is selective for flames and lights with glare, and suppressed by high SF black-and-white patterns including writing (Figure A19). These types of blur selectivity support categorical units in the final FC8 layer described below.

In the deeper FC layers, we examined units detected by our strict blur metric for size 155 and found that the vast majority of these units showed some form of blur selectivity. In FC7, three of the most prominent units based on our criterion were selective for entirely blurry or grainy images. For example, the preferred images for FC7 #4064 were blurry coffee makers or other objects (Figure A20). Another unit was selective for tigers or similar large cats (FC7 #3240, Figure A21), which have blur associated with the texture of animal fur and with the depth gradient in head-on photos.

In the final categorical layer, FC8, 36 of the 1000 categorical units were identified by our strict criterion based on responses to large shapes. The preferences of all 36 of these units, based on a subjective interpretation of their preferred and anti-preferred images, were related to blur. Some were selective for object categories that involved gradients in depth, like “steam locomotives” (unit #820, Figure A22) and “computer keyboard” (unit #508, Figure A23), where quotes are used to indicate category names. Some were selective for categories that are fundamentally blurry in appearance, such as “axolotl” (unit #029, Figure A24) and many were selective for animal fur, such as “Shetland sheepdog” (unit #230, Figure A25). Many were selective for rounded objects, such as “Granny Smith” apples (unit #840, Figure A26) and “hair spray” bottles (unit #585, Figure A27). Finally, some were selective for lighting flare, such as “lighter” (unit #626, Figure A28) and “stage” lights (#819, Figure A29). BCB units did not appear among the units identified by our strict criterion in FC8. For example, sharply focused birds or insects on blurry backgrounds were not among the preferred images; however, such patterns figured prominently among the anti-preferred images for the units detected by our criterion.

### Relative efficacy of simple shapes vs. natural images

The CNN tested here was trained only on natural images and not on simple, achromatic stimuli in our blurred shape set. To determine how well our shape set drove the CNN units relative to natural images, we averaged the maximum response to shapes across all units in each layer and divided this by the average maximum response to natural images. This ratio was higher in early layers, where units were often driven well by simple local combinations of orientation, boundary shape, luminance and spatial frequency, but diminished in deeper layers (Figure 7), where complex feature combinations (e.g. of color, texture and form) were likely required to drive these units but were not present in our shapes. In the version of AlexNet tested here, the Conv layers were divided into two sets of kernels in terms of allowable connections, originally for purposes of computational efficiency, that ended up causing differences in chromatic sensitivity. When we divide the Conv units into achromatic and chromatic sets (see Methods), it is apparent that the achromatic set is driven more strongly by our shapes (black line in Figure 7) whereas the chromatic kernels are driven less strongly (green line in Figure 7). In the FC layers, the distinction between chromatic and achromatic sets no longer exists at the architectural level, and we thus did not divide the units. In summary, our simple pre-defined shapes were able to drive the achromatic pathway at Conv2 to more than 33% of the average maximum achieved by searching through more than 36 million natural images. However, by FC7 and FC8, the shapes could not reach 10% of the average maximum responses to natural images. These numbers are consistent with results from related analysis for unblurred shapes (cf. Figure 13 in Pospisil et al., 2018). In spite of the substantial decline in the ability of the shapes to drive units deeper in the network, our strict blur criterion was still able to identify units that appear subjectively to be encoding categories of images containing blurry or low spatial frequency components.

**Figure 7.**
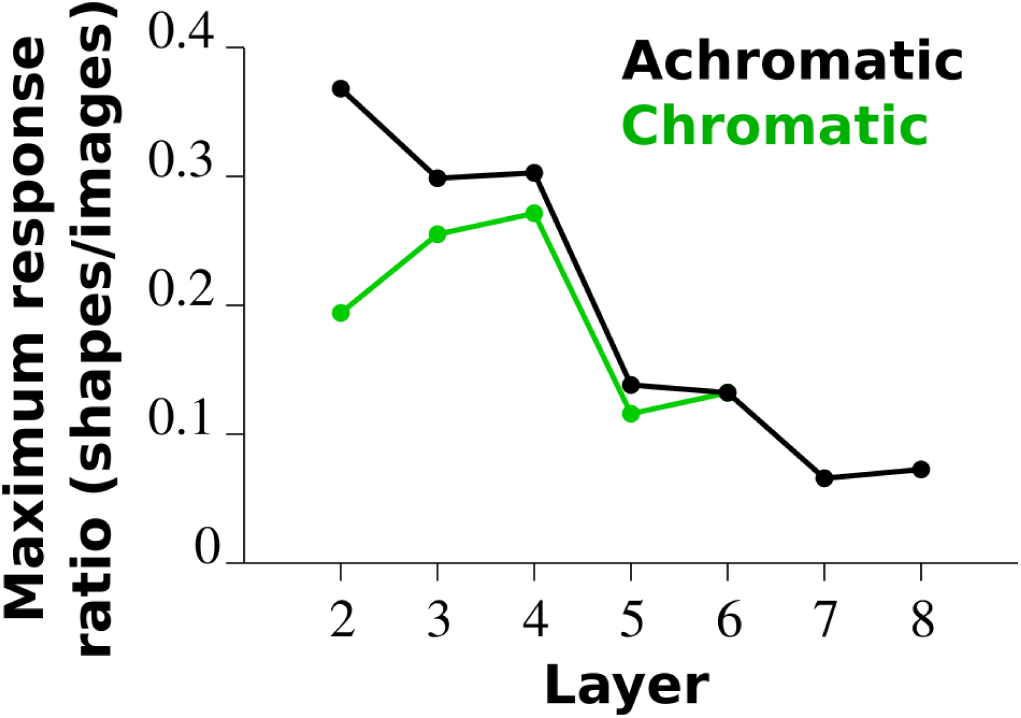
Maximal response to shape stimuli compared to that for natural images. The average (across units in a layer) of the maximum response (across all shapes and all blur levels, n=2,534) is divided by the same metric for 50,000 natural images (test set). This ratio of average maxima is plotted as a function of layer. For the Conv layers (2 to 5), the units are divided into an achromatic set (black line, half of all units in each layer) and a chromatic set (green line, the other half of units in each layer). For the FC layers (6 to 8), all units are averaged together. Number of units per layer is shown in Figure 3.

## DISCUSSION

To gain insight into the potential visual selectivity of units previously identified as encoding boundary blur in area V4 (Oleskiw et al., 2018), we adopted the visual stimulus set from the macaque experiments to identify, within a CNN, blur selective units from shallow to deep layers that could then be studied in depth. Major conclusions from our study are as follows: (1) there are a substantial fraction of units in the CNN that appear to be involved in blur encoding when tested with simple, artificial stimuli, (2) several subjective forms of blur selectivity emerged, including BCB units that are selective for boundaries defined by blur contrast, (3) blur selectivity is transferred and transformed across layers, (4) units selective for blank or relatively uniform image patches are also detected by our simple blur assay, and (5) many units involved in blur encoding are missed by our assay. Our results call for in vivo experiments to determine whether BCB units are prominent in vivo, particularly in area V4. They also suggest that more refined stimulus sets are needed to quantify and characterize blur selectivity in vivo and in artificial visual systems.

### Forms of blur selectivity

Our subjective classification of blur selective units includes the following. Blur region units are responsive to blurry regions covering their entire RF and may be broadly tuned for orientation. BCB units are selective for oriented or shaped boundaries between regions containing different levels of blur, e.g., different spatial frequency content. These BCB units may be thought of as textural analogs to double-opponent color neurons in macaque visual cortex (for review, see Shapley and Hawken, 2011). Instead of chromatic and spatial opponency, BCB units have a form of textural and spatial opponency. Other units respond best to low frequency oriented gradients or transitions between relatively blank regions defined by luminance or color. In future work, we aim to develop quantitative criterion to distinguish these classes.

### Shallow vs. deep units

In Conv2, our blur criterion detected many units that were selective for boundaries between high SF and low SF regions, e.g., between a sharply focused bird or insect and a blurry background. Such units may be important not only for finding object boundaries, but also for determining the location of the object to be classified within an image. Such boundaries are low-level features that may be relevant to many types of objects and visual imagery. It is not surprising that at the deepest, categorical level (FC8) our assay did not detect such units. In the deepest layers, the units identified as blur selective corresponded to categories associated with rounded forms, like bottles or apples, or with extended soft regions lacking sharp object boundaries, like animal fur, or with objects that have prominent depth gradients in photographs (steam engines and keyboards). Blur selectivity may be fundamentally different at the low and high level. At the low level, it relates to differences in spatial frequency, whereas at a high level, real blur is distinct from blank or homogenous regions (Ma et al., 2018).

### The use of simple artificial stimuli

For decades there has been an ongoing debate about the use of simple artificial vs. natural stimuli in the biological vision literature. Our approach directly tests the ability of simple stimuli to accurately detect units that have a desired visual selectivity. In this case, a simple assay based on blurry shapes was sufficient to identify many potentially blur selective units that also appeared to be selective for blurry natural stimuli. On the other hand, the use of achromatic or monochromatic stimuli could fail to detect certain types of blur-selective units. In particular, many units were exclusively suppressed by our stimulus set (e.g., Figure 2C). It would be important to use a stimulus set that explores color in future assessments of blur selectivity. In addition, our stimuli are not ideal for detecting BCB units, because these units are both excited and suppressed by blur. A stimulus set that explicitly includes blur-defined boundaries needs to be developed that is effective for characterizing these units.

### Relationship to neurophysiology

Our study was motivated by a previous report of boundary blur encoding in V4, and our results are consistent with those in V4 in terms of the approximate proportions of blur-selective units detected - roughly on the order of 20% of units (Oleskiw et al., 2018). To determine whether blur-encoding neurons in V4 map onto the subjective classes defined here may require experiments that use far more stimuli than possible given practical limitations. It is also possible that earlier visual areas may have units with blur selectivity similar to what is reported here. For example, units that detect second-order (not luminance-defined) texture boundaries have been found and studied in the cat visual cortex (Mareschal and Baker, 1998; Baker and Mareschal, 2001). Their finding that the orientation of the inputs to such units may bear no relationship to the orientation of the overall boundary detected by the unit is consistent with our results. In particular, our visualization of BCB units shows that many different orientations may be involved in driving the units. Few studies have examined texture-defined form in macaque visual cortex, but one major study concluded that V2 is likely not directly involved in encoding texture-defined form (El Shamayleh and Movshon, 2011). In the future, it will be important to test a greater variety of stimuli in V2, and to pursue such units downstream in area V4.

To improve our assay of blur-selectivity, it would be useful to apply an independent quantitative metric of blur. For example, a measure of relative power at low vs. high spatial frequency in the preferred vs. anti-preferred images could be used to independently identify units that are blur selective. There are also image blur metrics available from computer vision, including a very simple difference between neighboring pixels summed along all horizontal and vertical lines that has been shown to successfully identify out of focus image regions (Li et al., 2001). However, to properly detect BCB units, such metrics would need to be constrained to particular regions of the image, thus complicating the procedure. Such metrics may be of limited use for identifying blur selective neurons in vivo, where one typically is unable to carry out a brute-force assessment of the best and worst images.

Future studies are now required to address several aspects of blur selective units in artificial neural networks: (1) what are the underlying circuits that create blur selectivity, (2) how are the outputs integrated with information in deeper layers to solve segmentation and classification tasks and (3) what sets of stimuli would be most effective for quickly identifying and quantitatively distinguishing classes of blur selectivity? The answers to the latter question can be worked out in networks and then applied in vivo, where a major open question involves understanding the encoding of texture-defined boundaries along the V1 to V2 to V4 pathway.

## ACKNOWLEDGEMENTS

This work was funded by NIH R01-EY027023 and R01-EY029997. We thank Anitha Pasupathy for helpful advice.

## APPENDIX

For example CNN units referred to in the main text, below we present the top five (excitatory) and bottom five (suppressive) images (with faces blocked by black boxes), and the deconv-net visualizations for the Conv layers, where maximal RFs are smaller than the entire input image. We have examined the top and bottom ten images for each unit, and find that the first five are always representative. For FC layers, the potential RF is the entire image and the deconv-net visualization does not provide great insight, thus only preferred and anti-preferred images are shown (top & bottom 10). The following list indicates the figures to follow, one per page.

**Figure A1.**
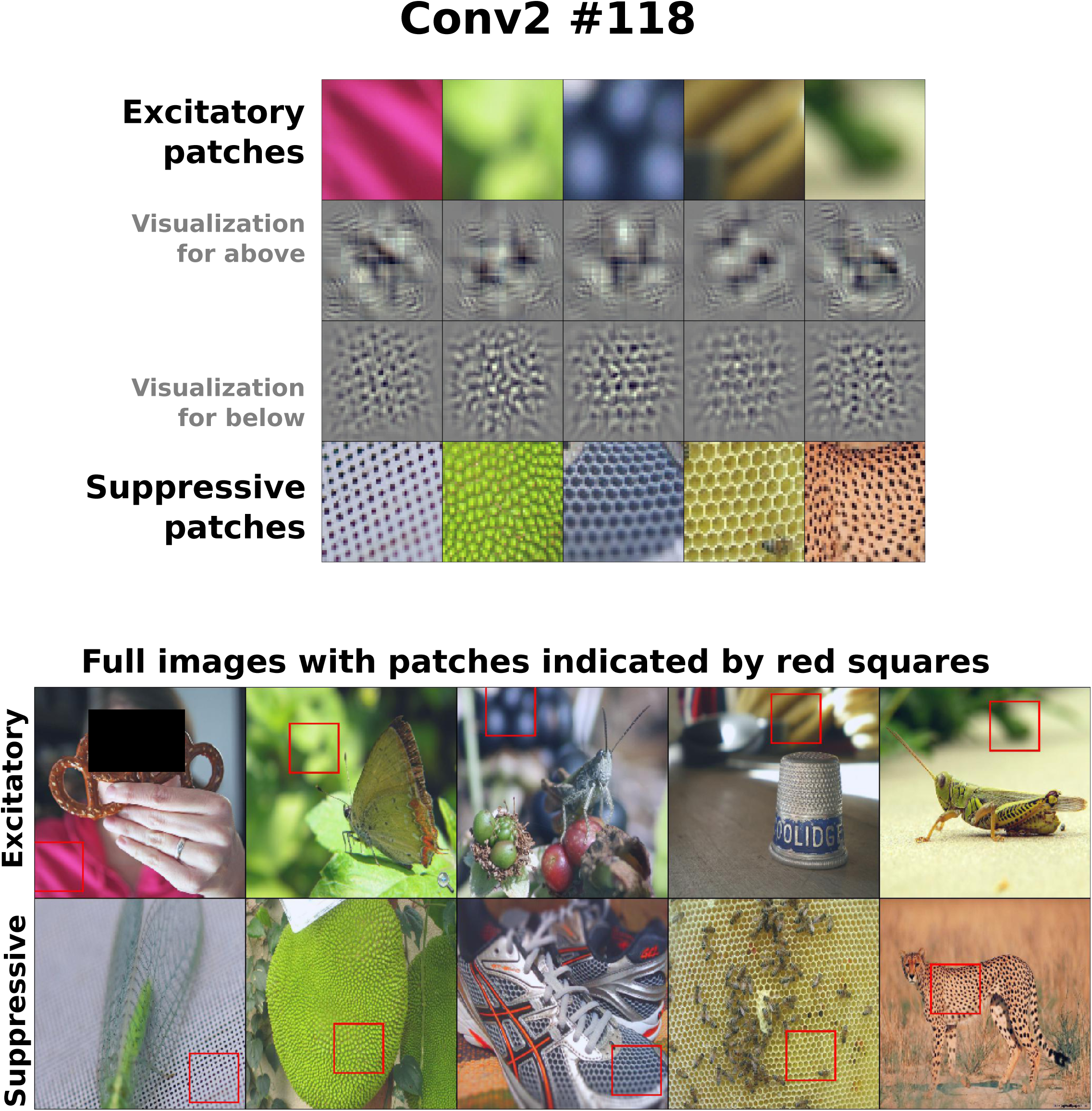
Conv2 #118 - example unit in Figure 2A.

**Figure A2.**
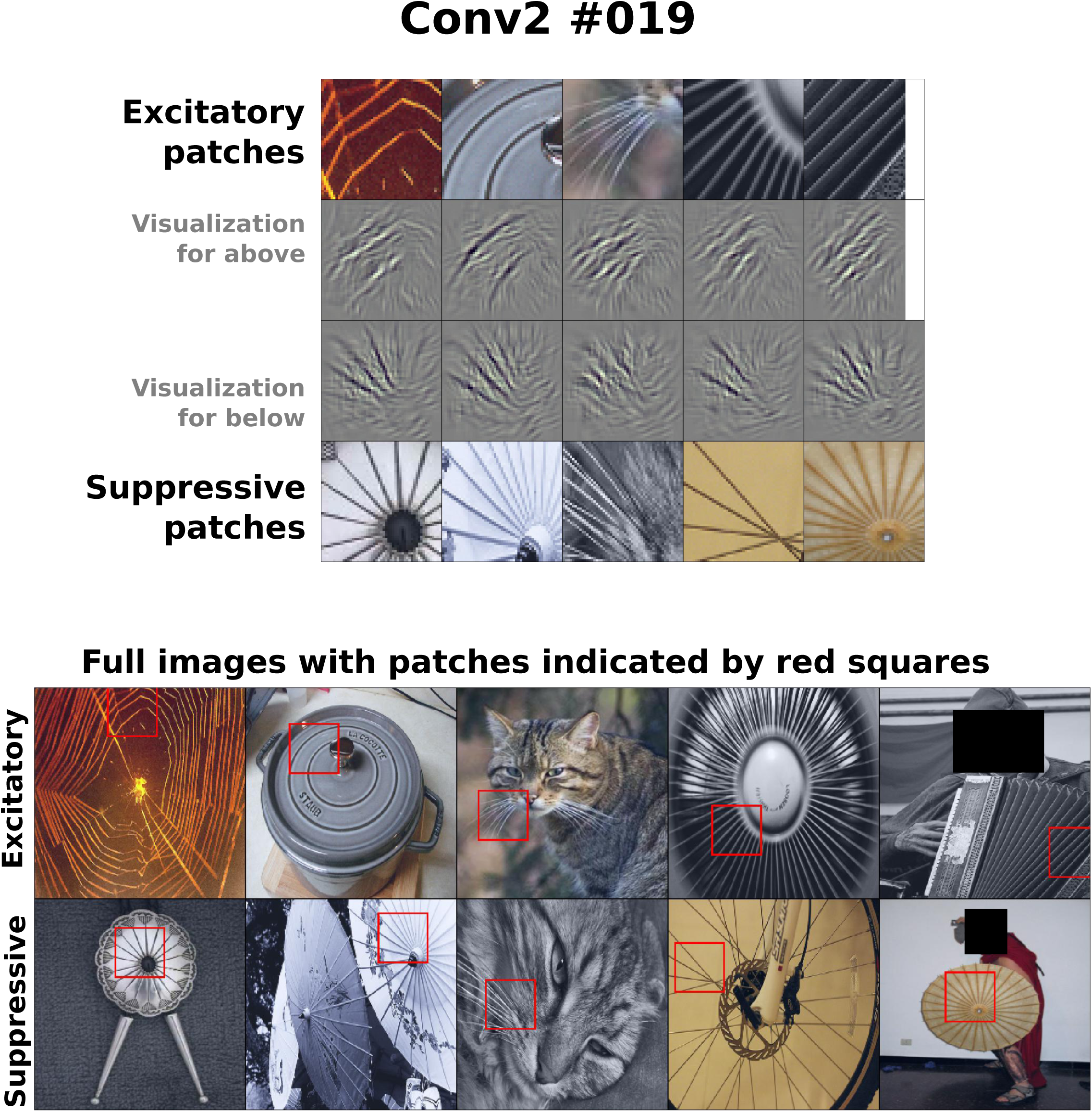
Conv2 #019 - example unit in Figure 2B.

**Figure A3.**
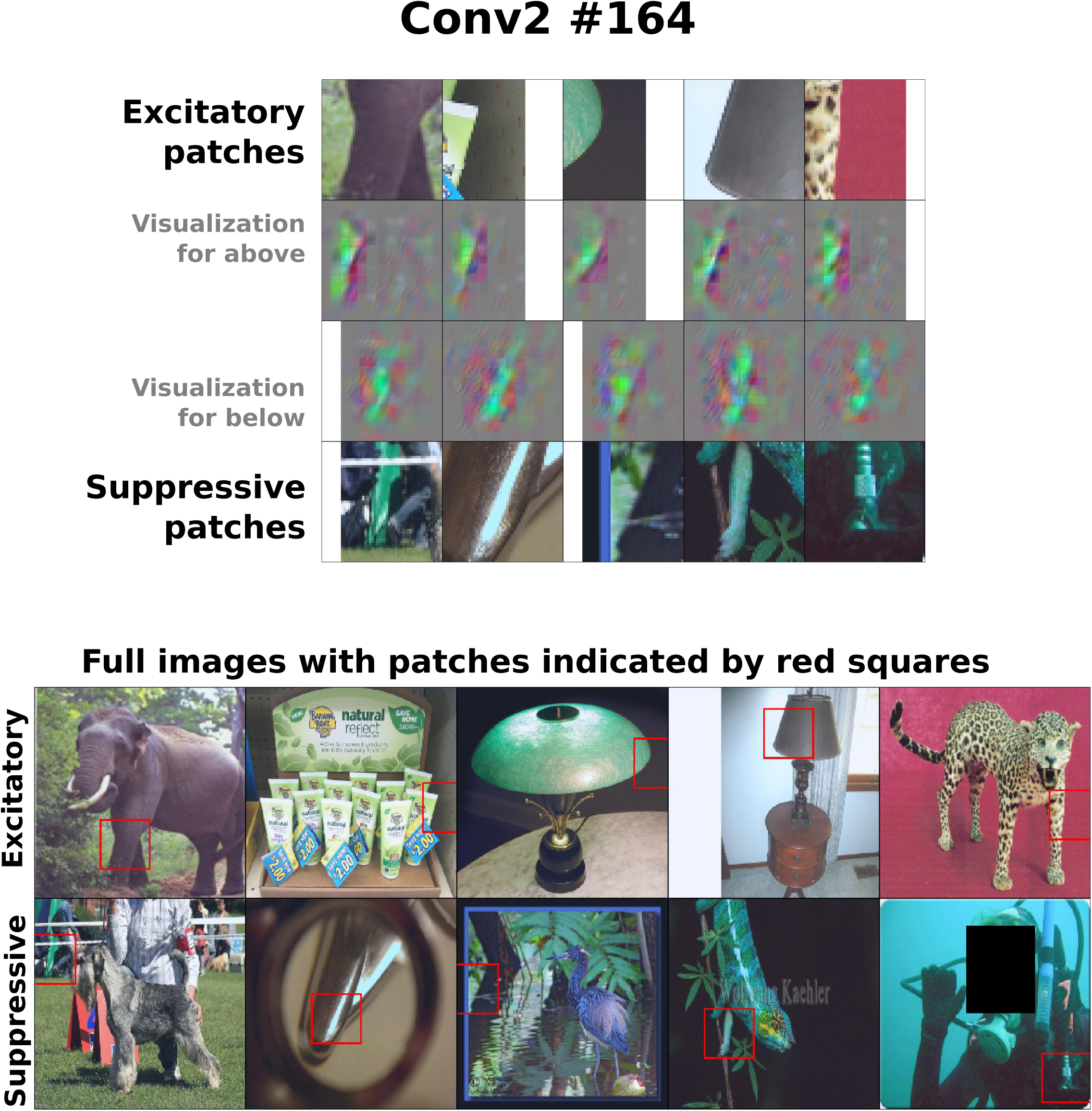
Conv2 #164 - example unit in Figure 2C.

**Figure A4.**
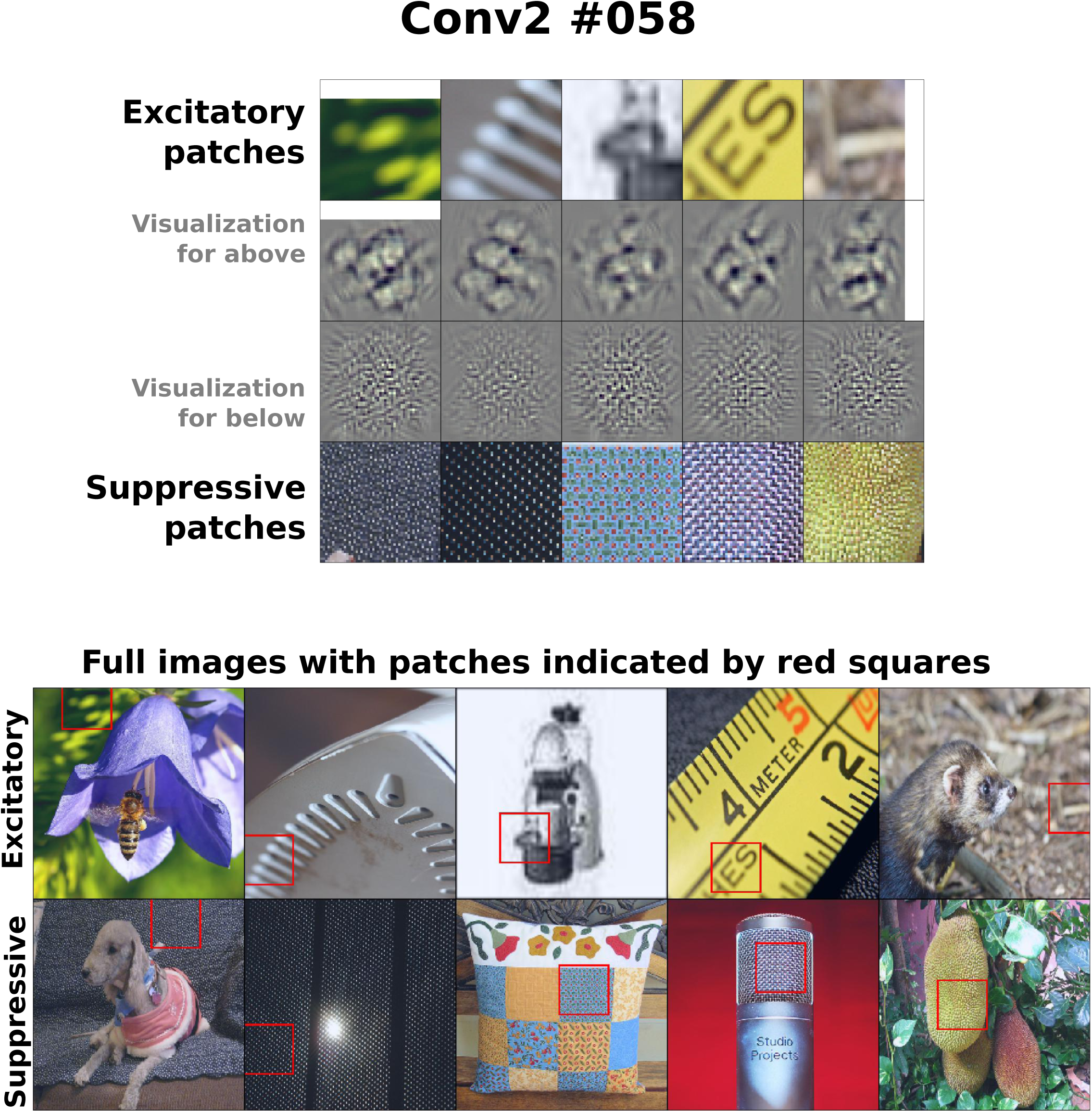
Conv2 #058 - second point from right in Figure 5A.

**Figure A5.**
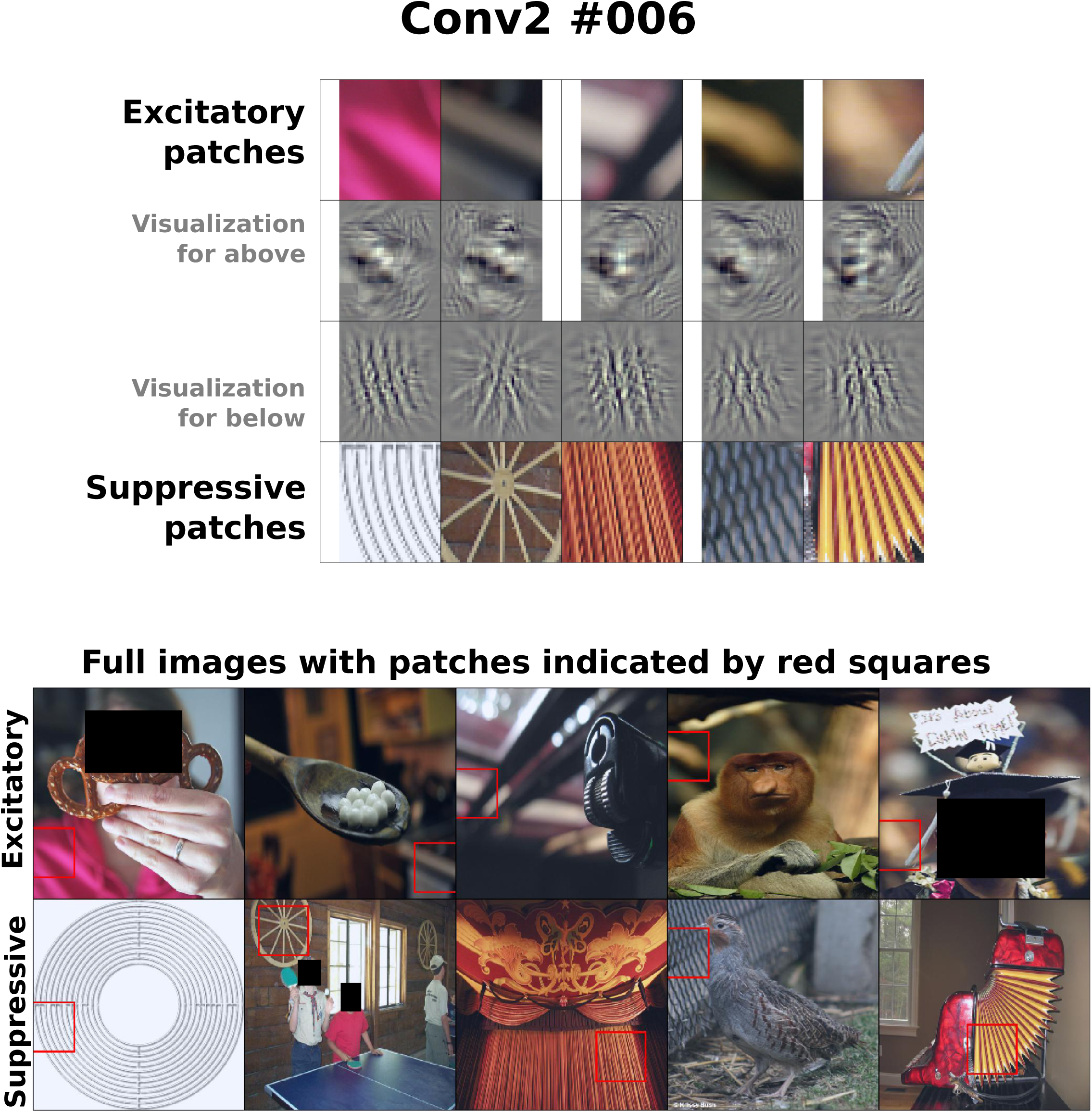
Conv2 #006 - third point from right in Figure 5A.

**Figure A6.**
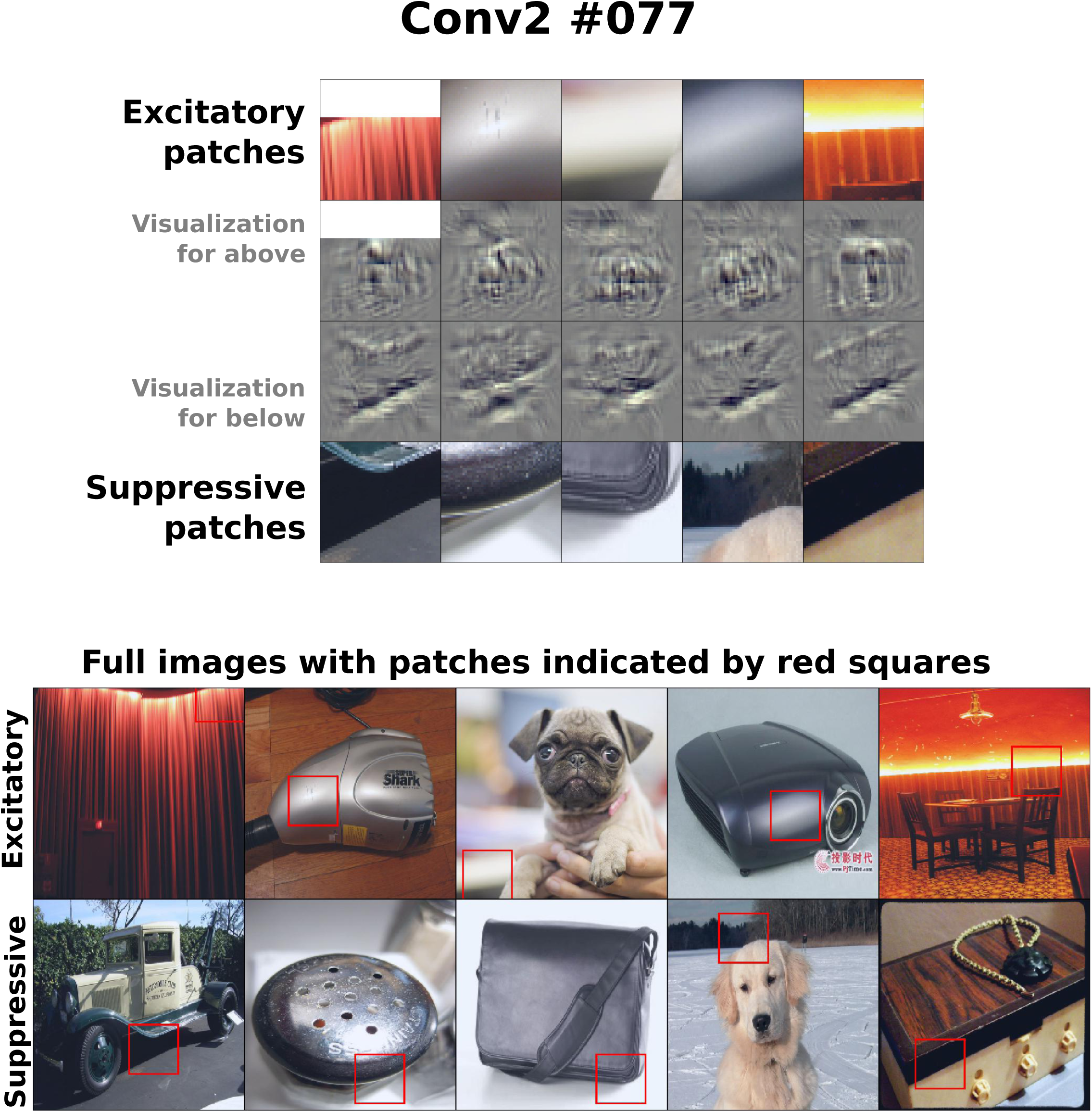
Conv2 #077 - example gradient unit.

**Figure A7.**
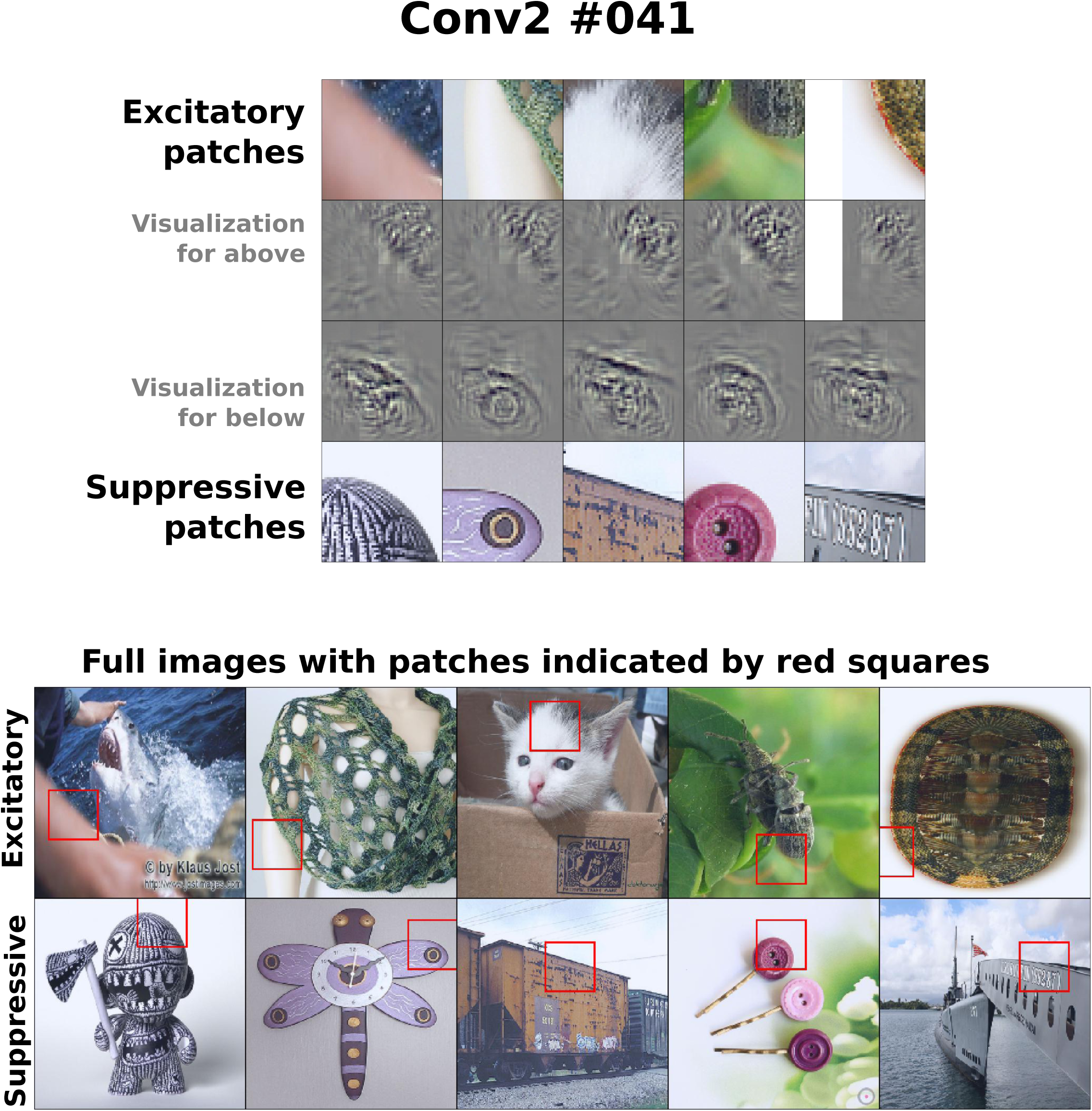
Conv2 #041 - a blur-contrast boundary (BCB) unit identified by strict criterion.

**Figure A8.**
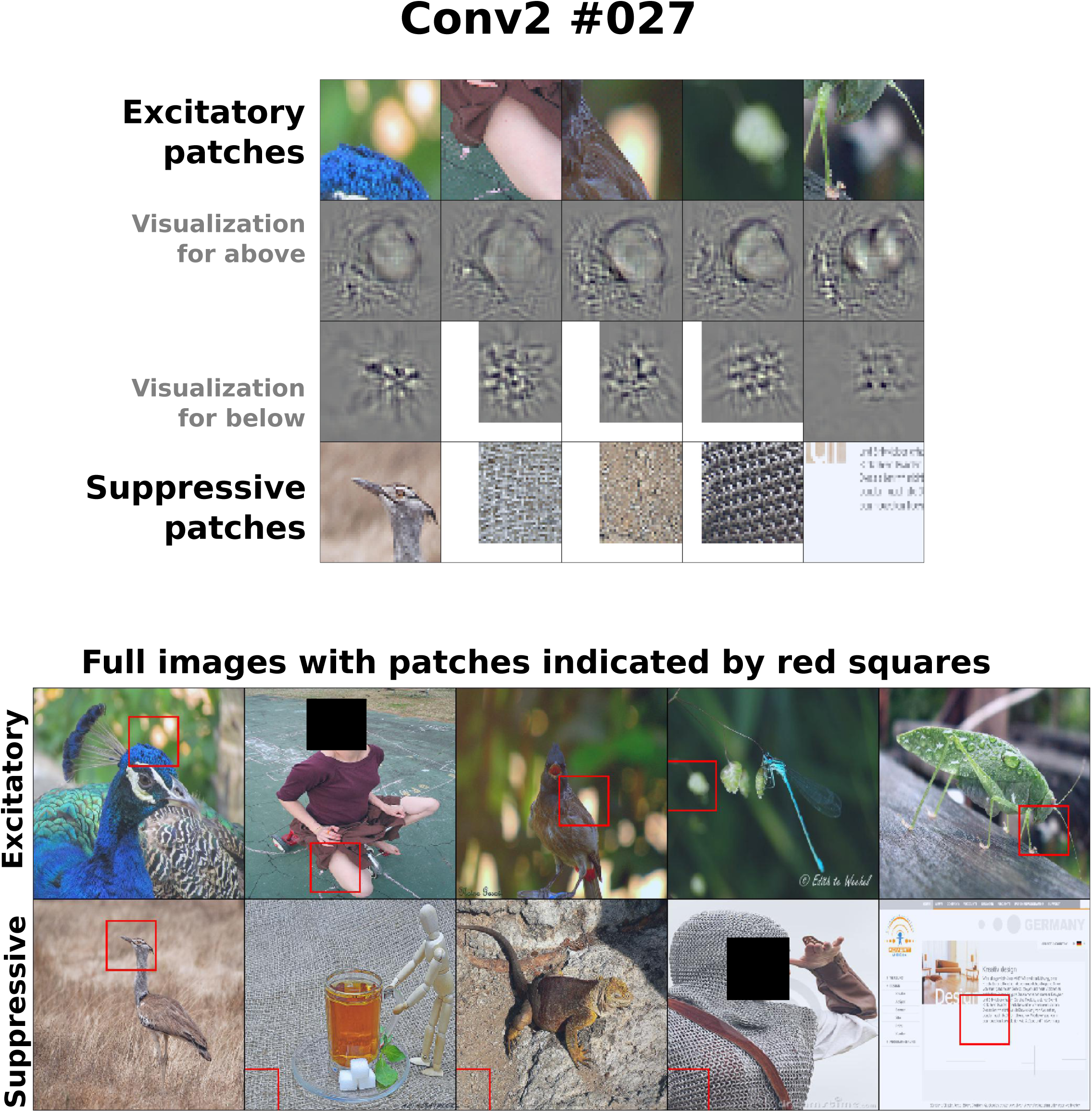
Conv2 #027 - a blur-contrast boundary (BCB) unit identified by lax criterion.

**Figure A9.**
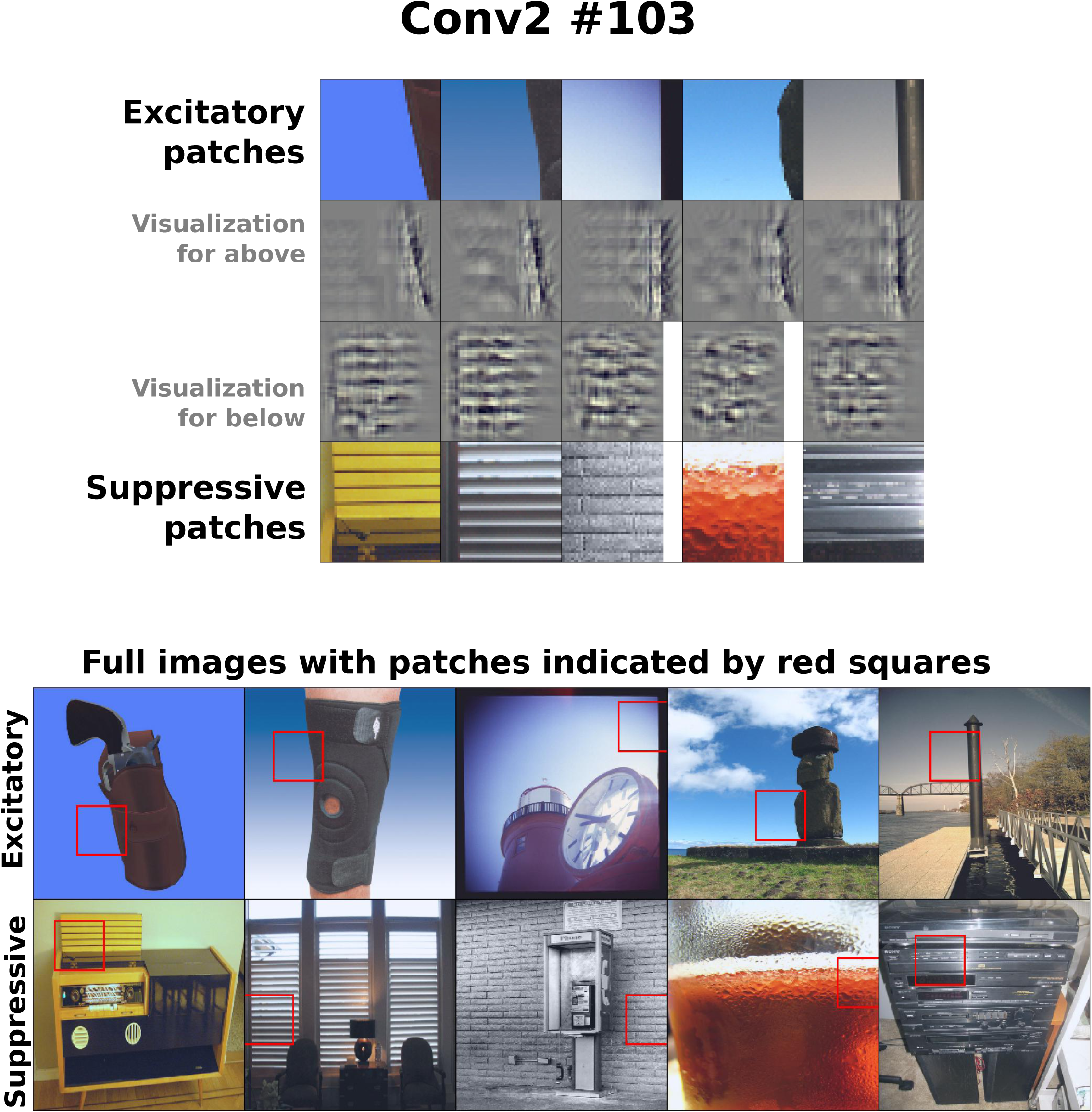
Conv2 #103 - a blur-contrast boundary (BCB) unit identified by lax criterion.

**Figure A10.**
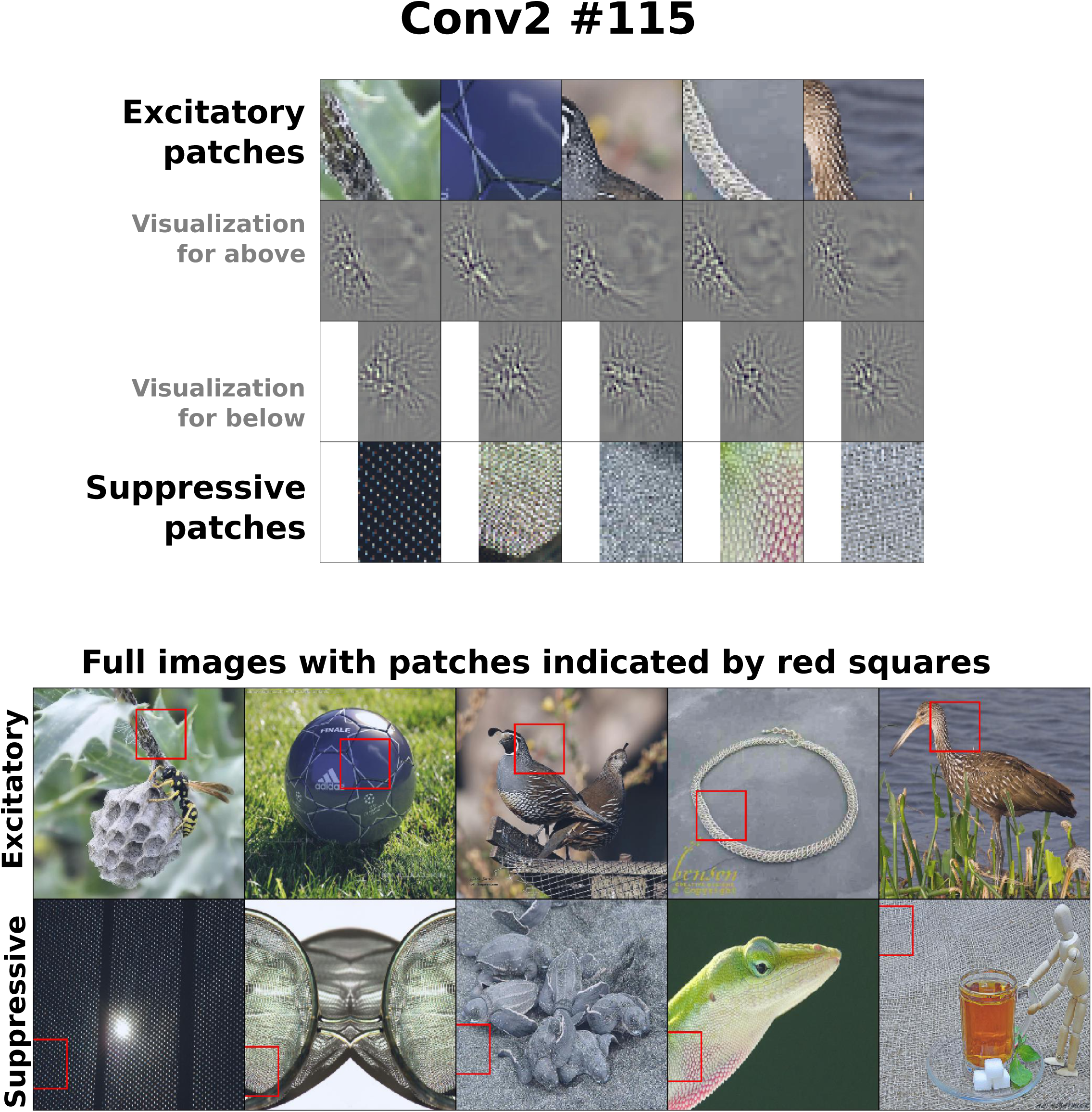
Conv2 #115 - a blur-contrast boundary (BCB) missed by strict and lax criteria.

**Figure A11.**
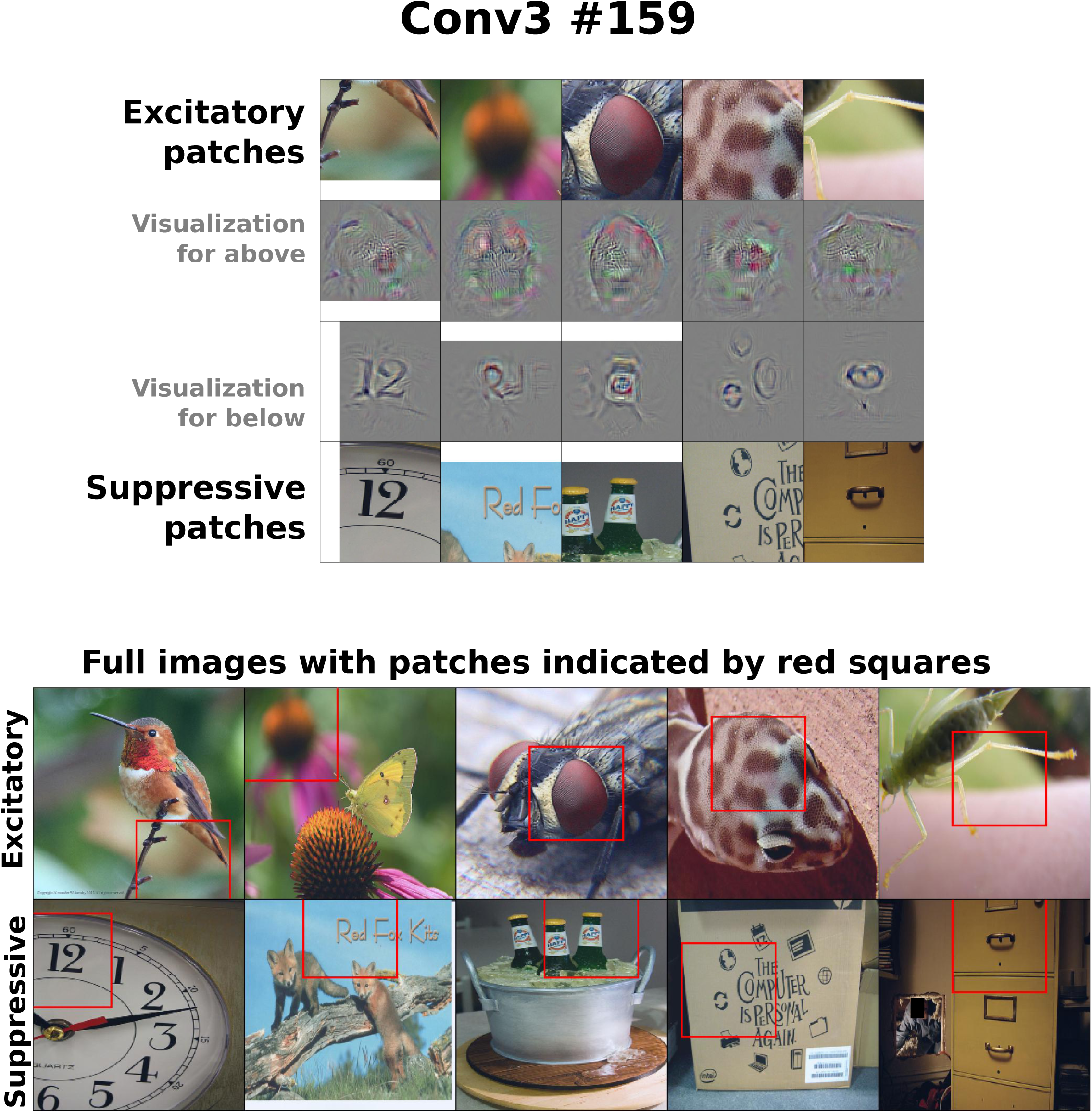
Conv3 #159 - right-most point in Figure 5B.

**Figure A12.**
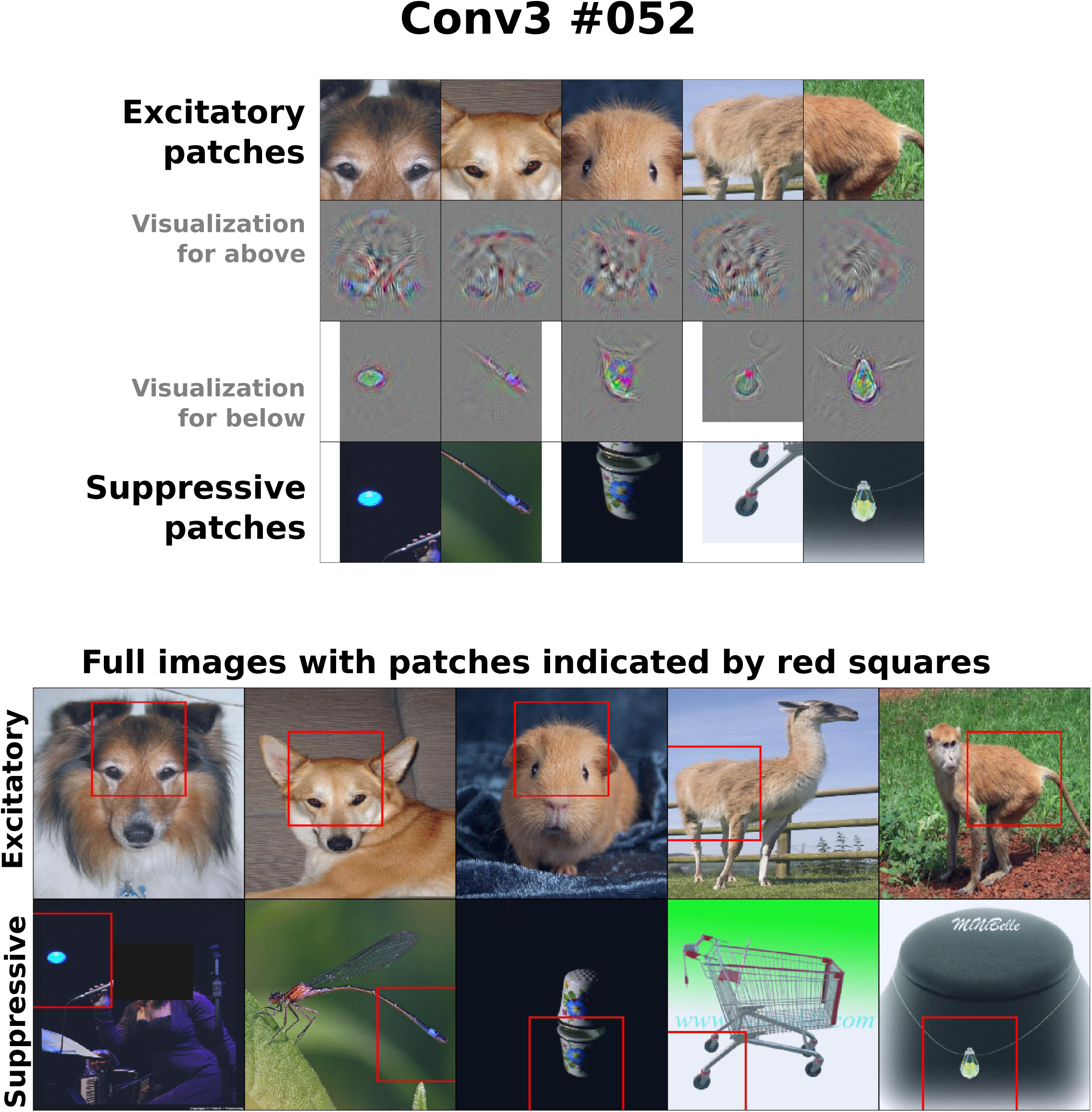
Conv3 #052 - second point from right in Figure 5B.

**Figure A13.**
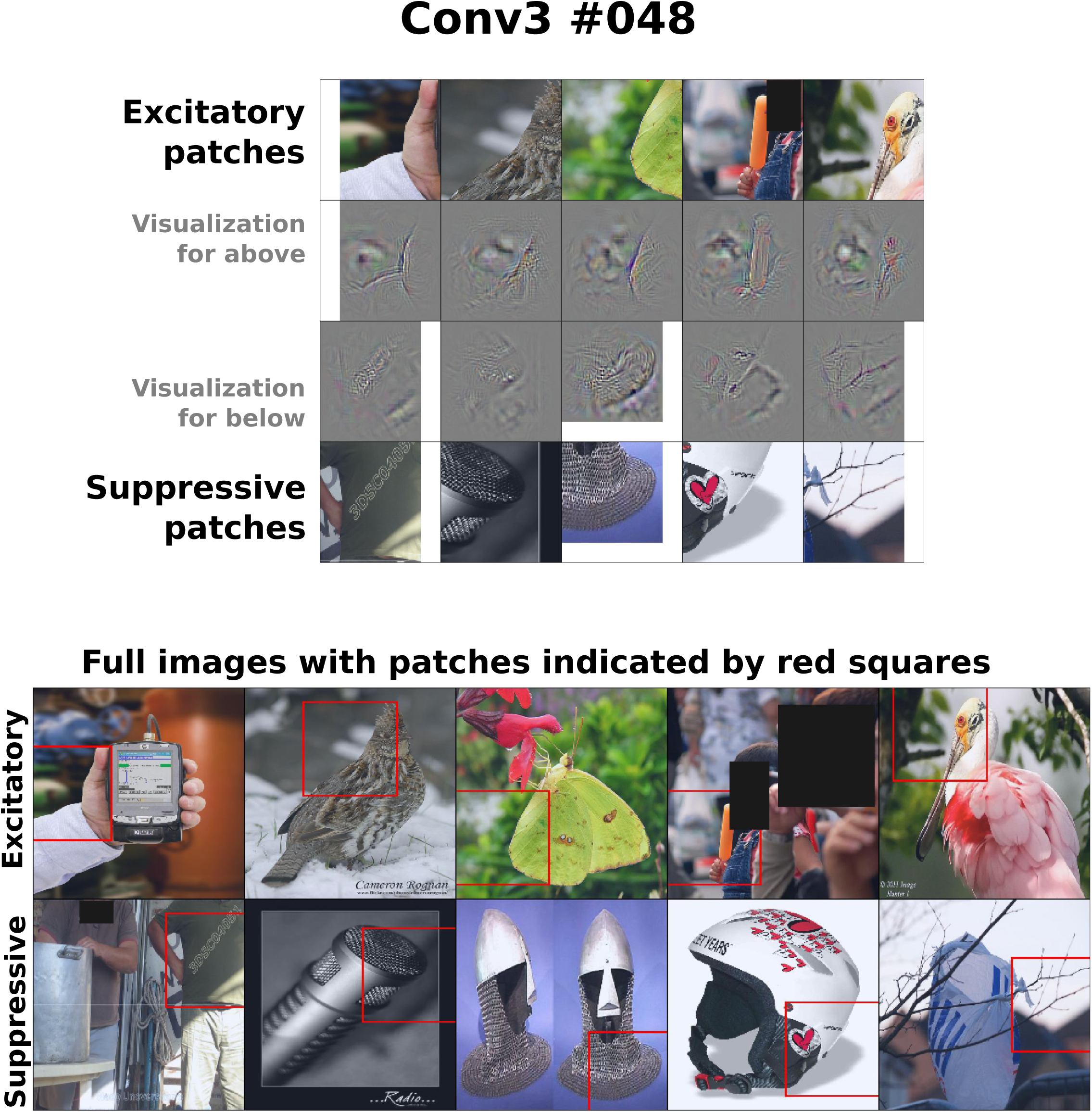
Conv3 #048 - BCB unit identified by strict criterion.

**Figure A14.**
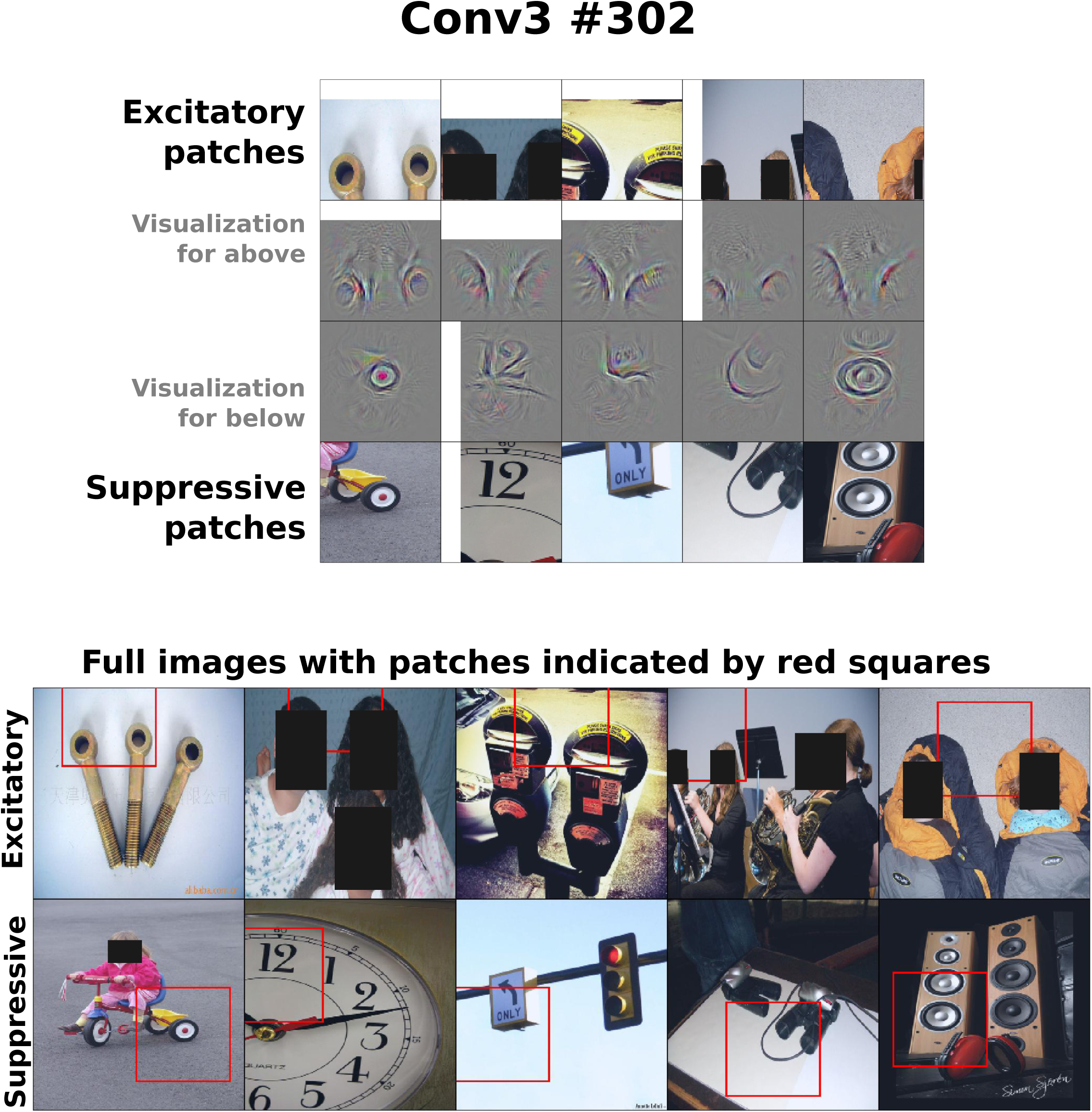
Conv3 #302 - fourth point from right in Figure 5B.

**Figure A15.**
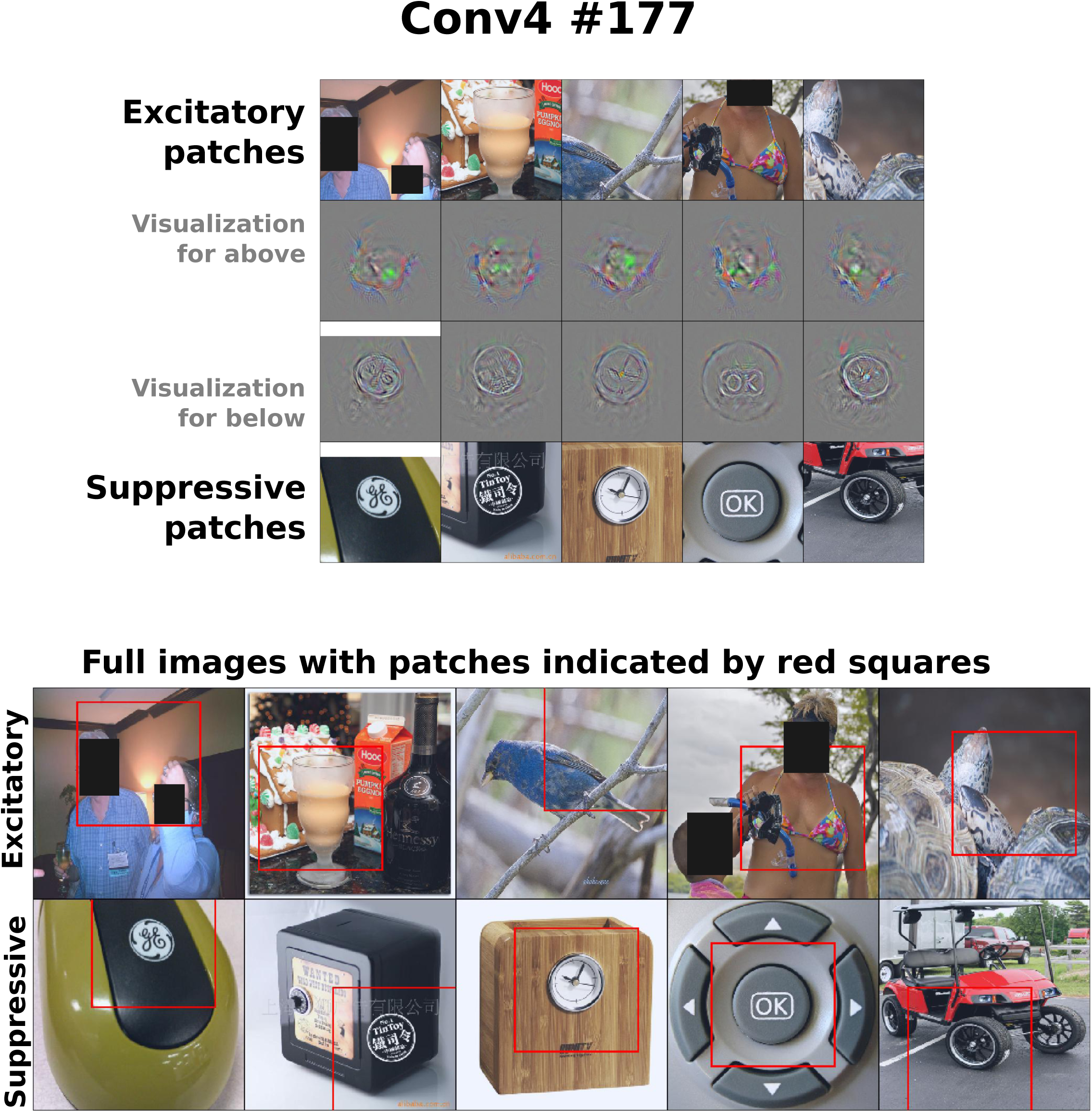
Conv4 #177 - right-most point in Figure 5C.

**Figure A16.**
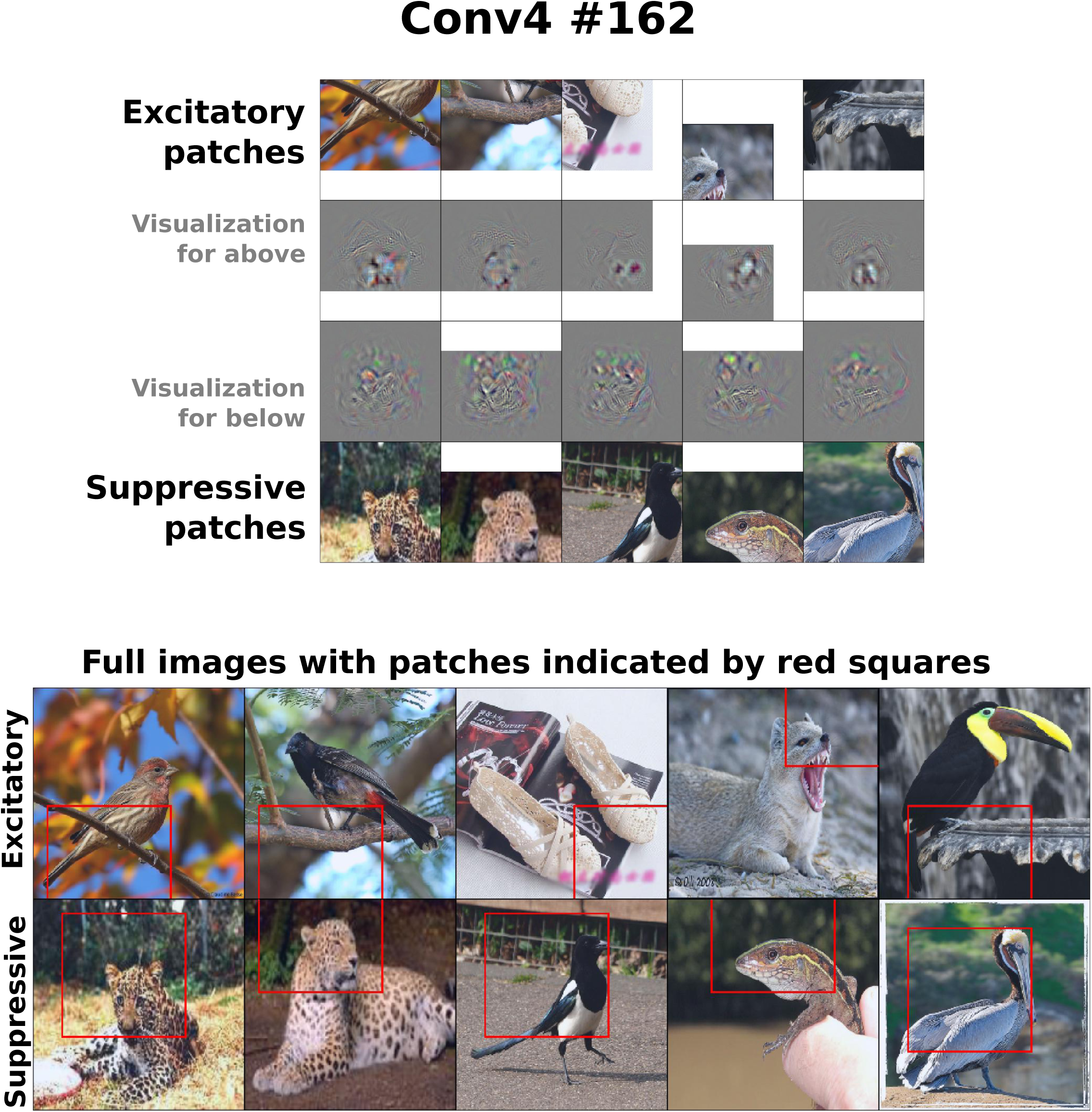
Conv4 #162 - third point from right in Figure 5C.

**Figure A17.**
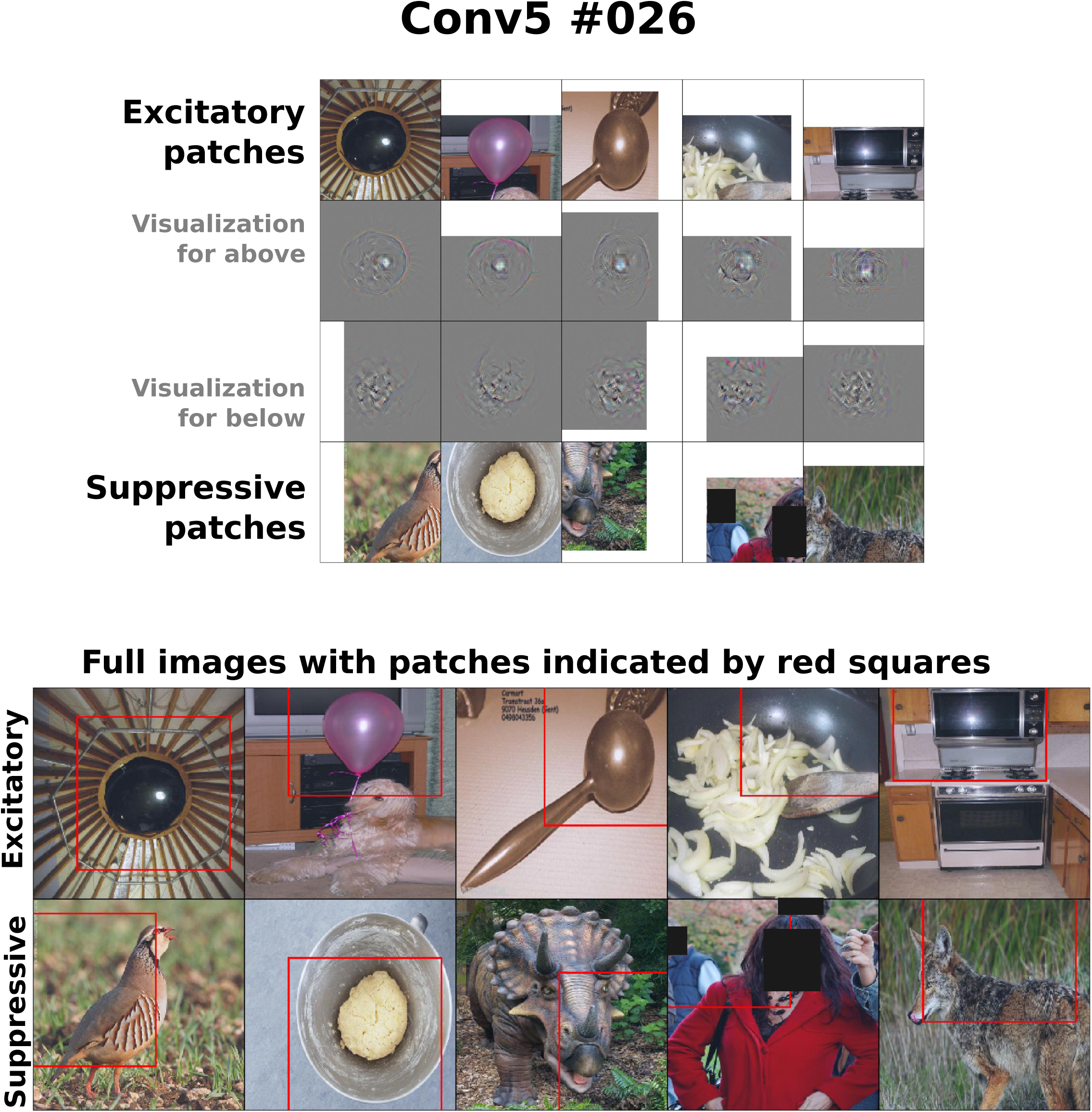
Conv5 #026 - right-most point in Figure 5D.

**Figure A18.**
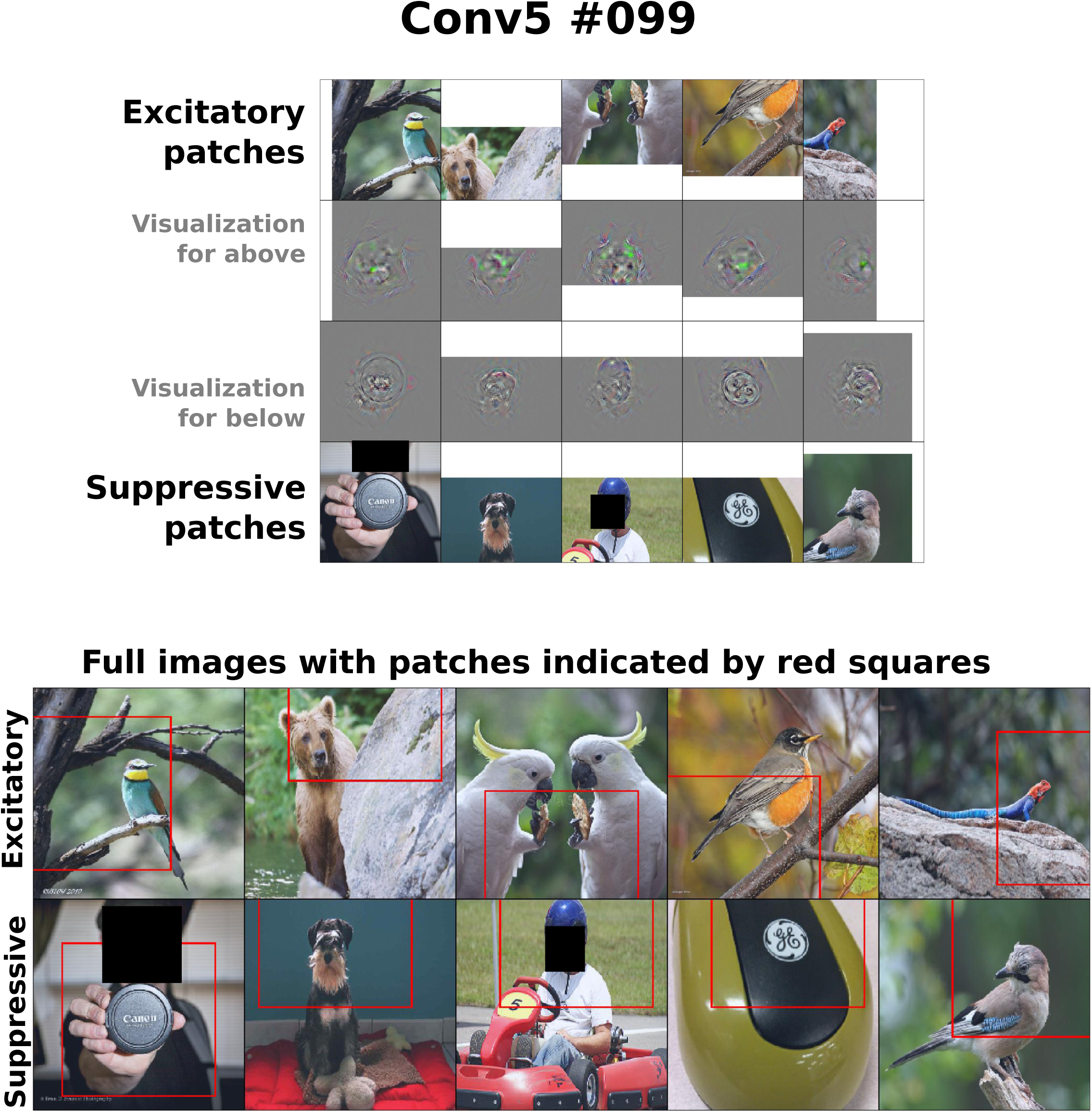
Conv5 #099 - third point from right in Figure 5D.

**Figure A19.**
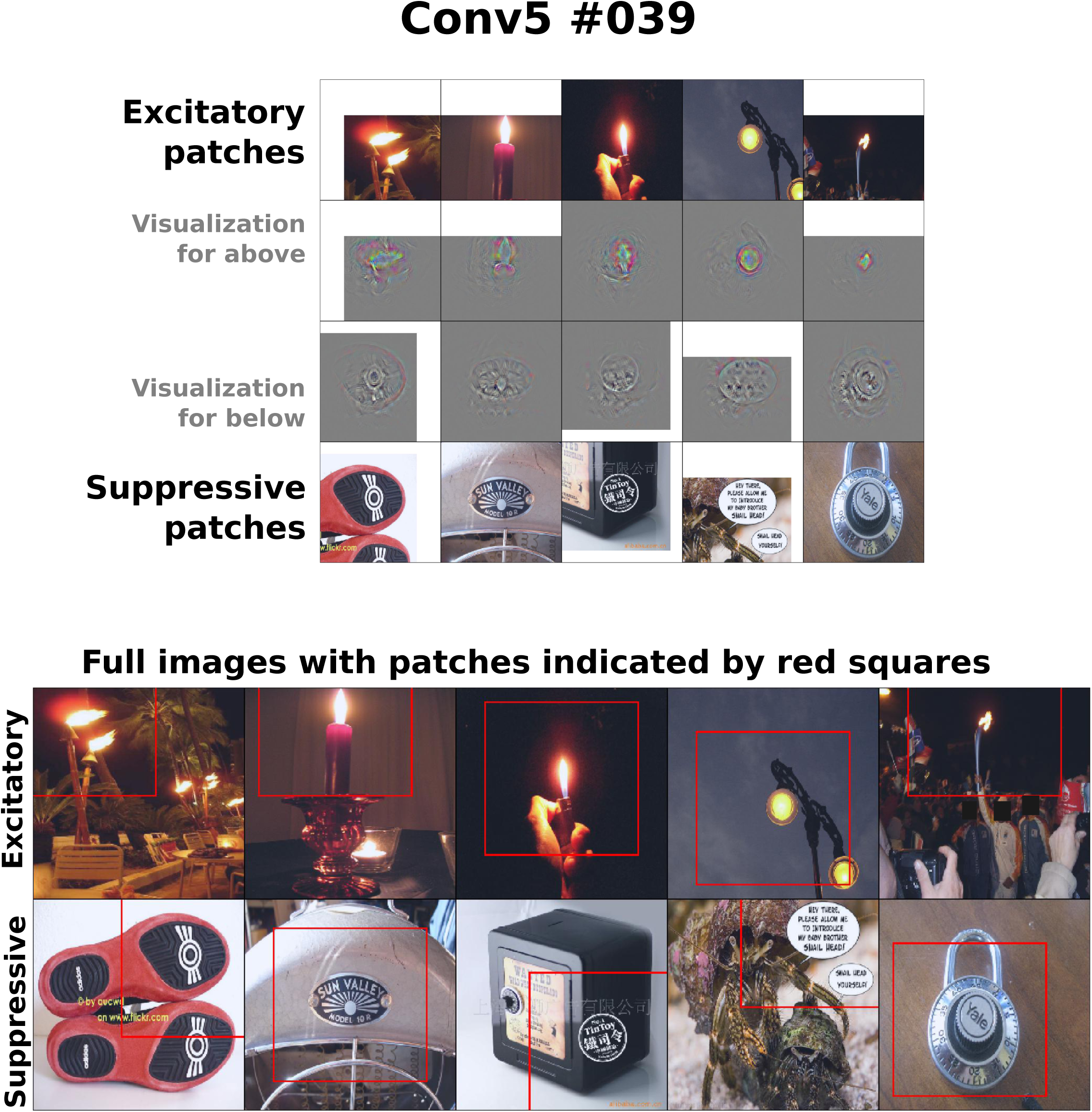
Conv5 #039 - example unit selective for a flame.

**Figure A20.**
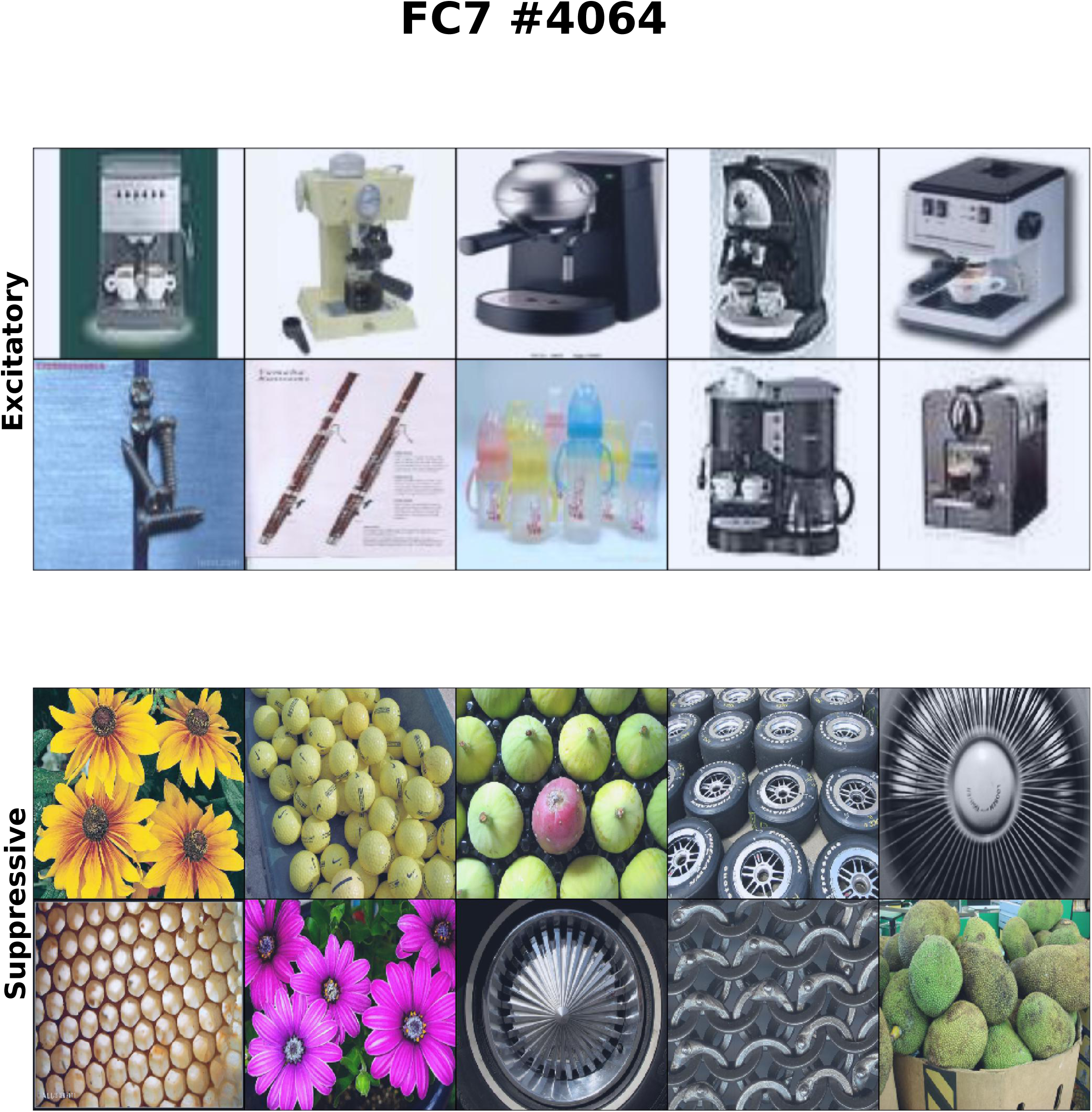
FC7 #4064 - example unit selective for blurry images.

**Figure A21.**
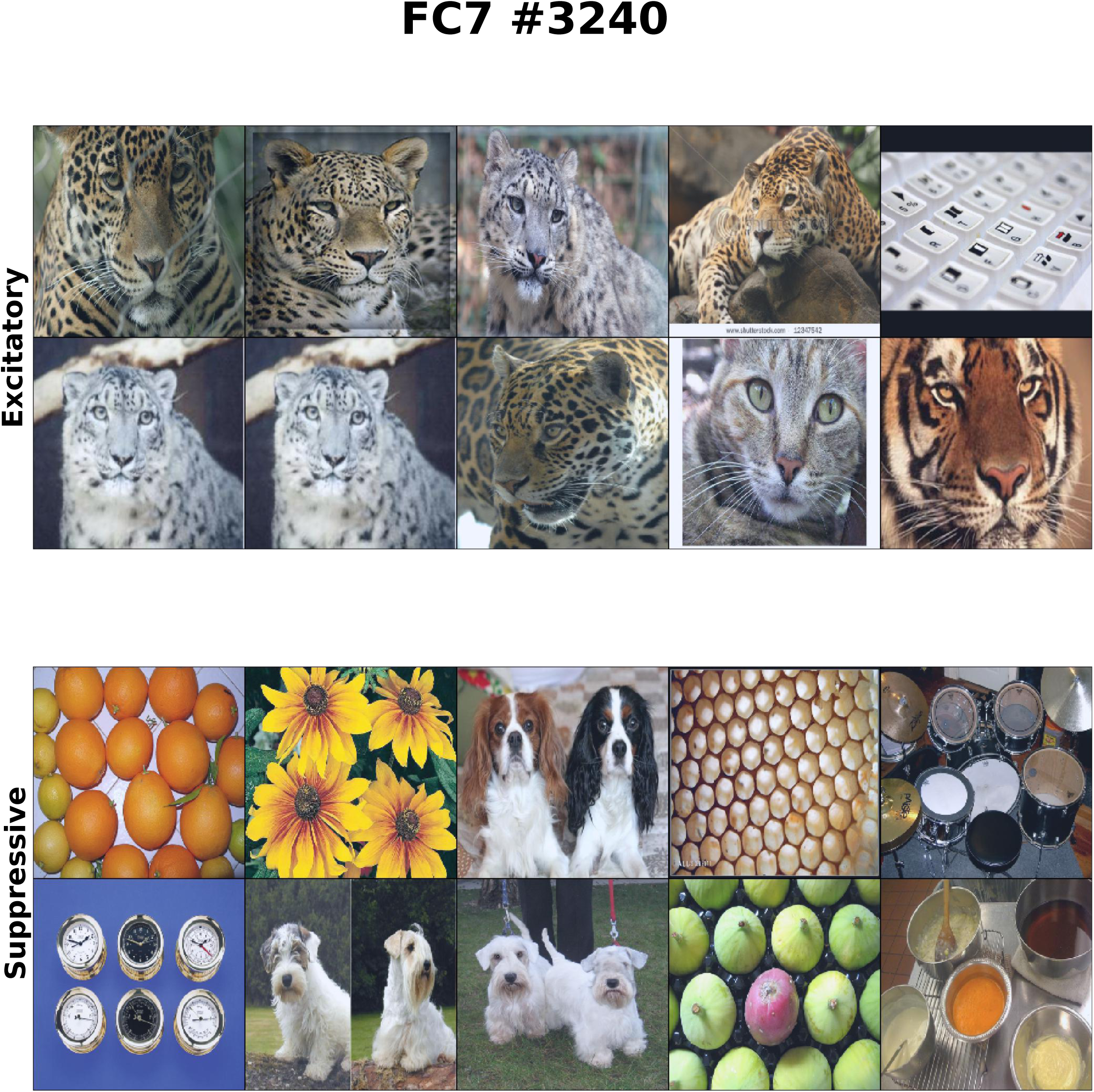
FC7 #3240 - example unit selective for animal fur.

**Figure A22.**
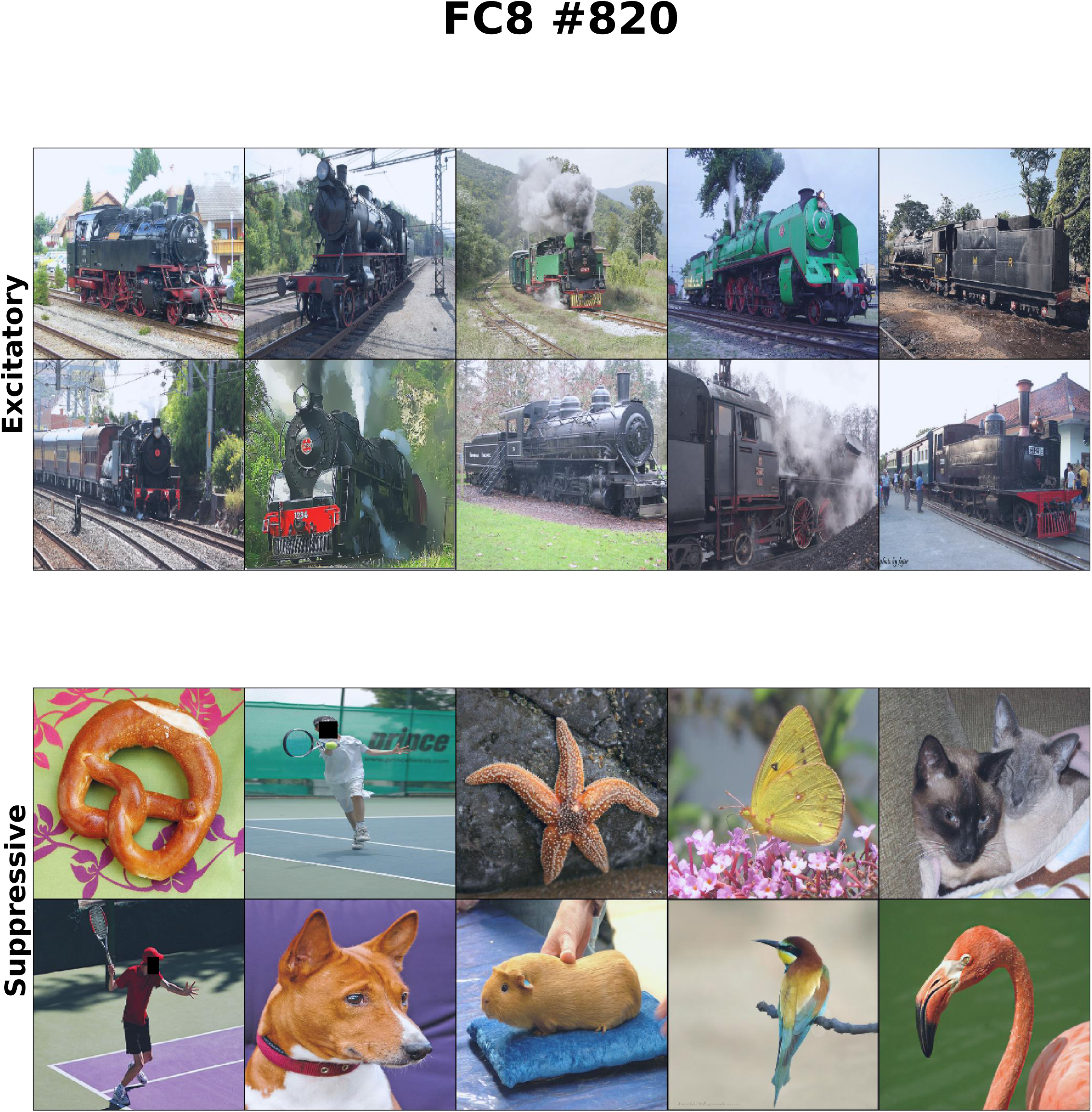
FC8 #820 - example unit related to depth gradient. Category: steam locomotive.

**Figure A23.**
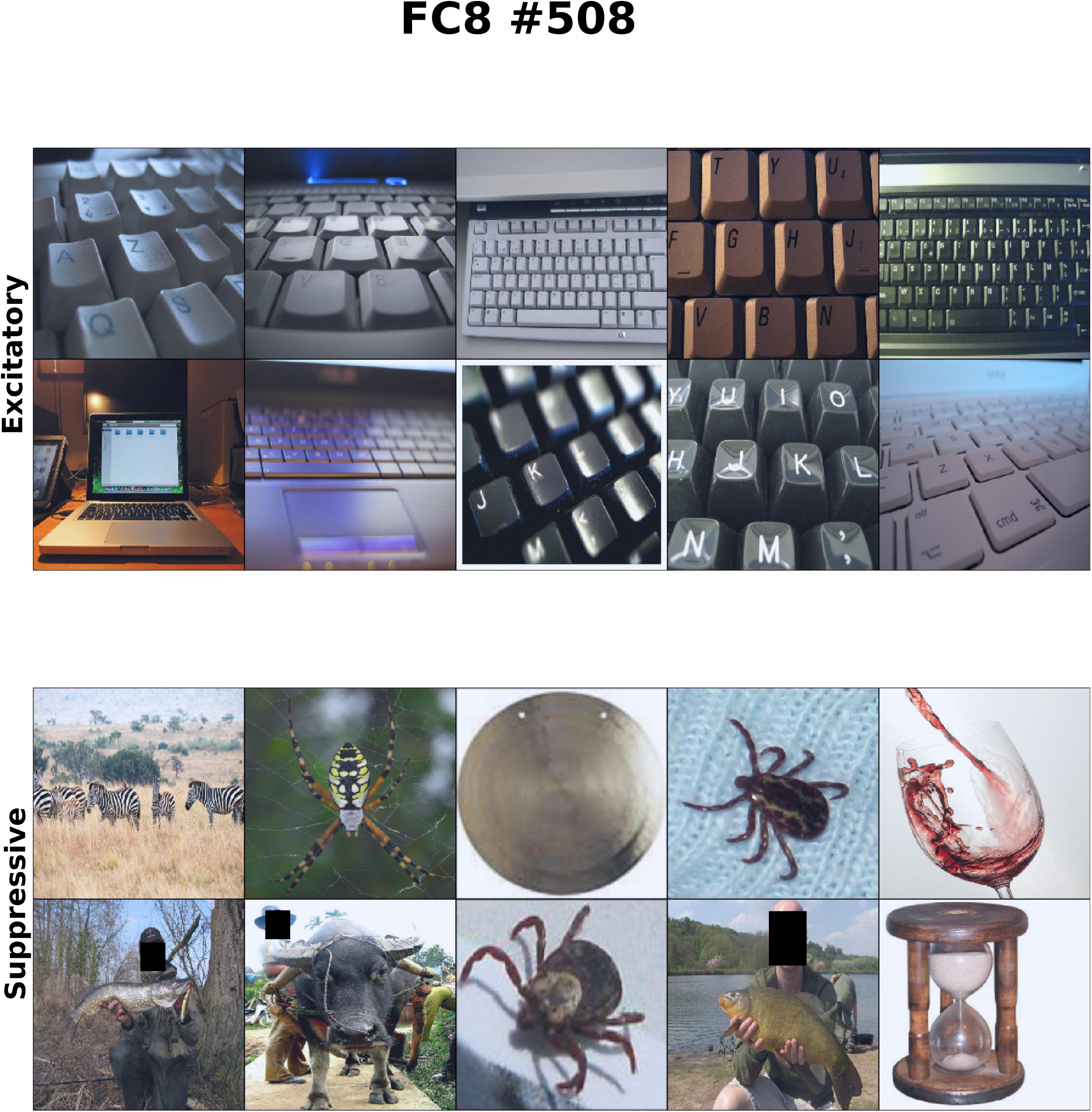
FC8 #508 - example unit related to depth gradient. Category: computer keyboard.

**Figure A24.**
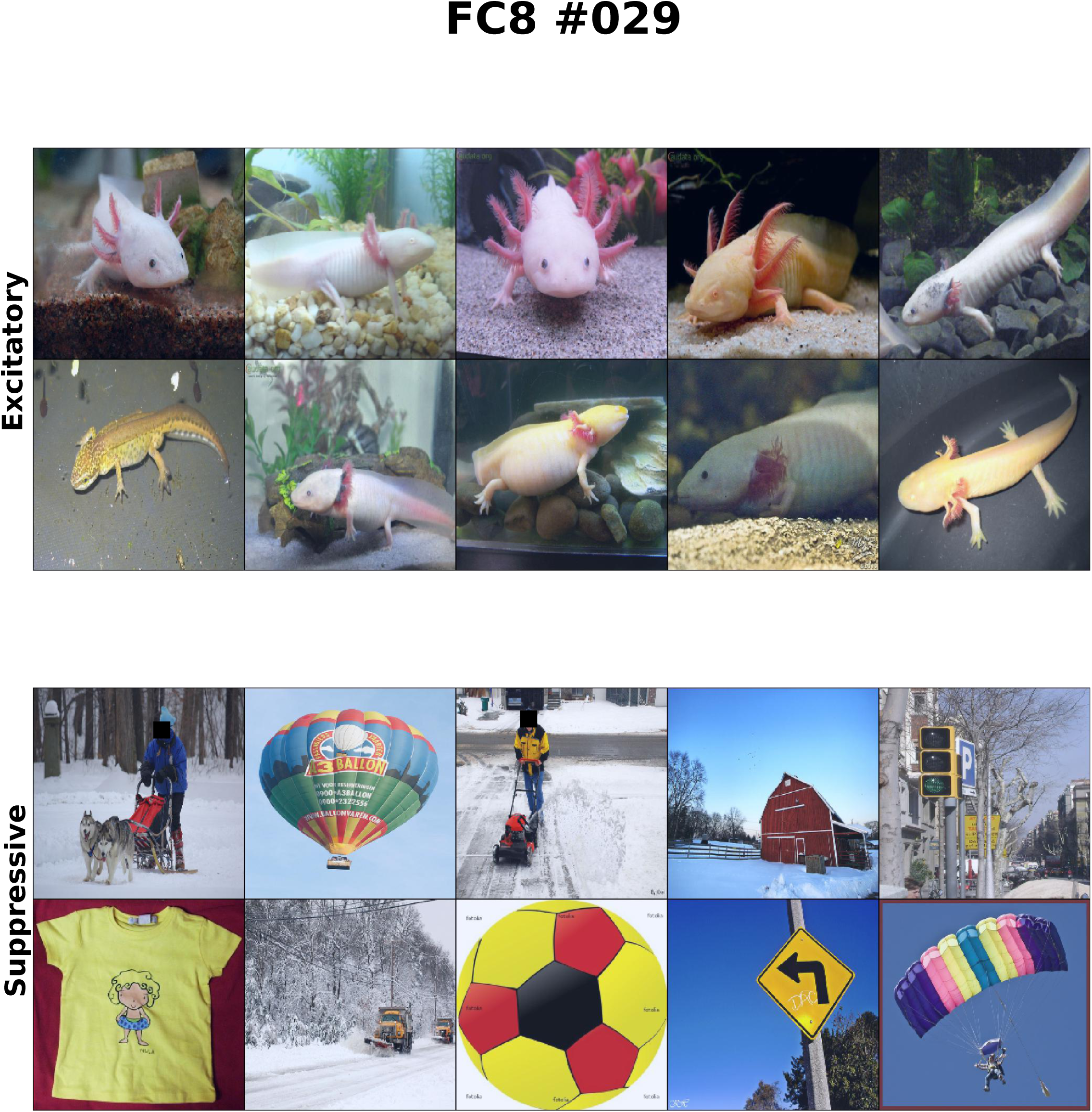
FC8 #029 - example unit selective for a blurry type of object. Category: axolotl.

**Figure A25.**
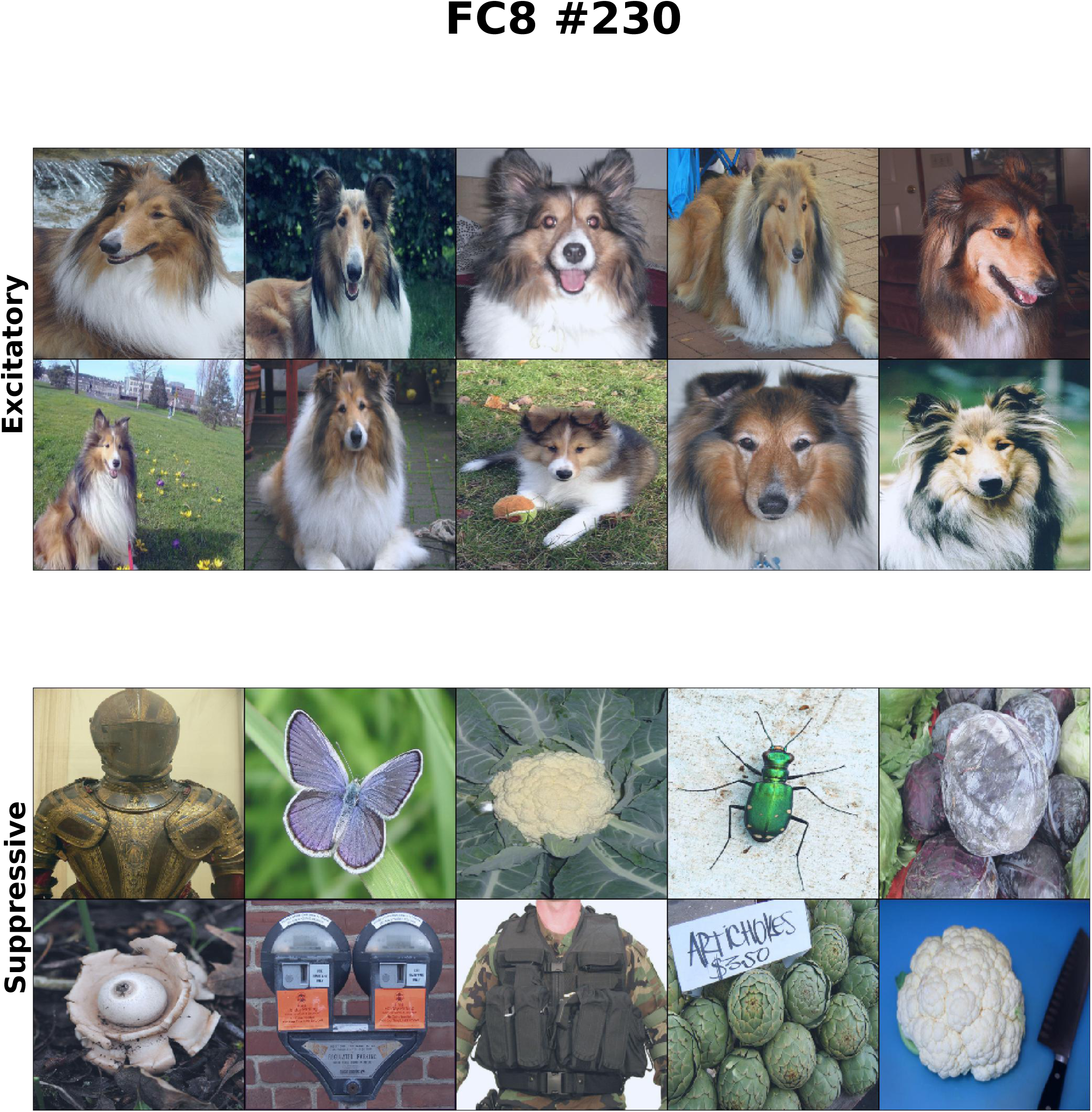
FC8 #230 - example unit selective for animal fur. Category: Shetland sheepdog.

**Figure A26.**
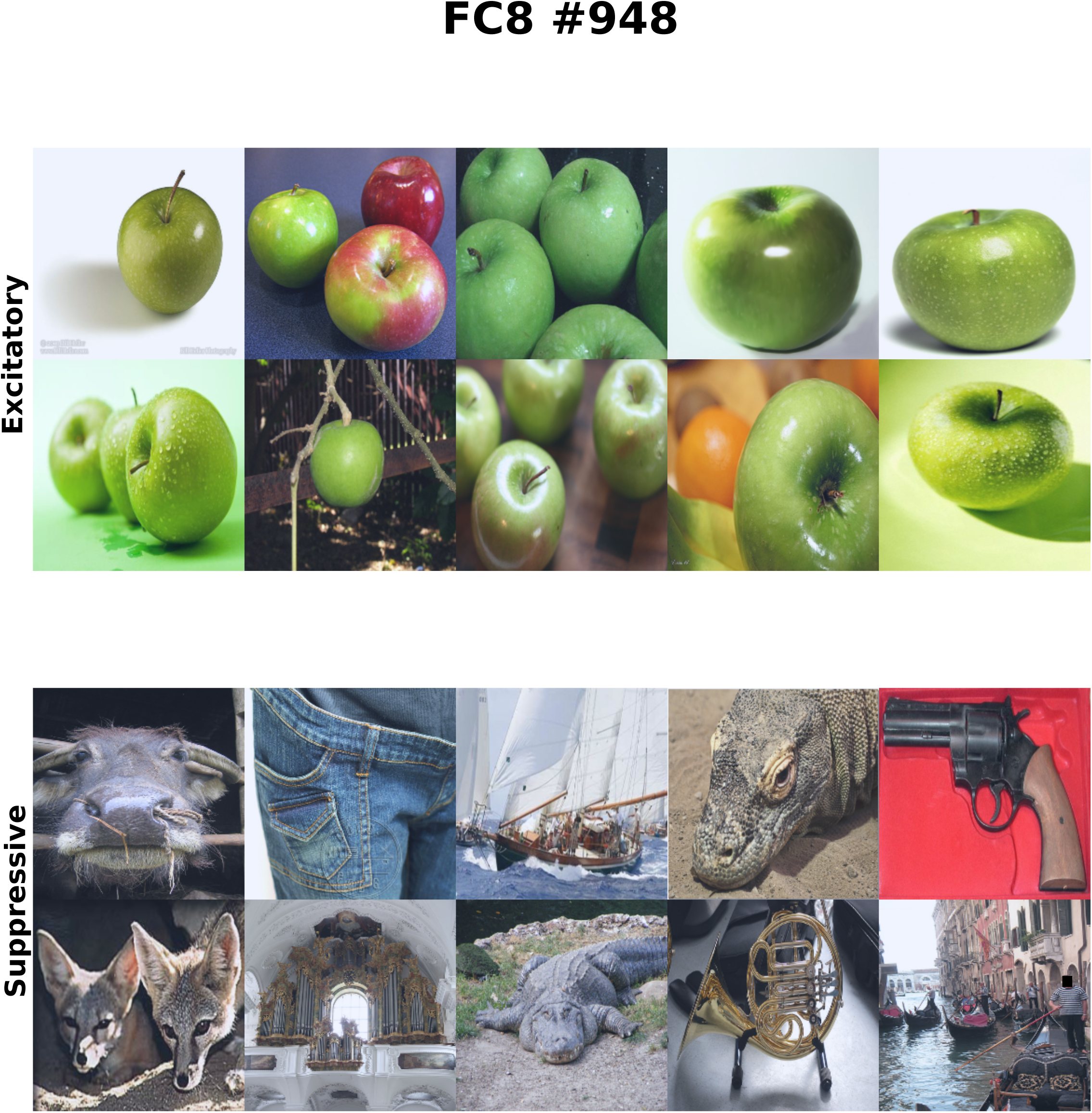
FC8 #948 - example unit selective for rounded objects. Category: Granny Smith.

**Figure A27.**
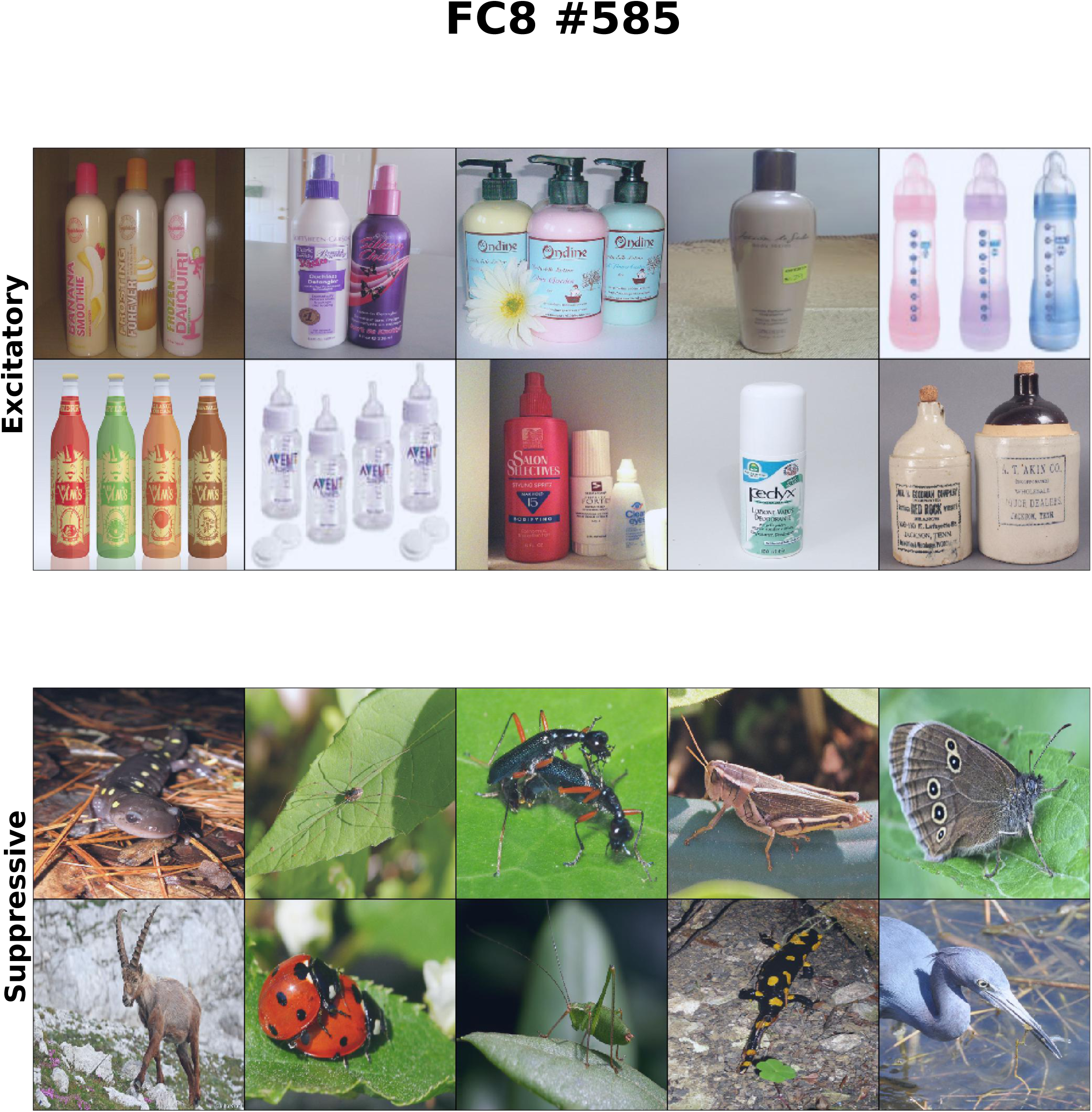
FC8 #585 - example unit selective for rounded objects. Category: hair spray.

**Figure A28.**
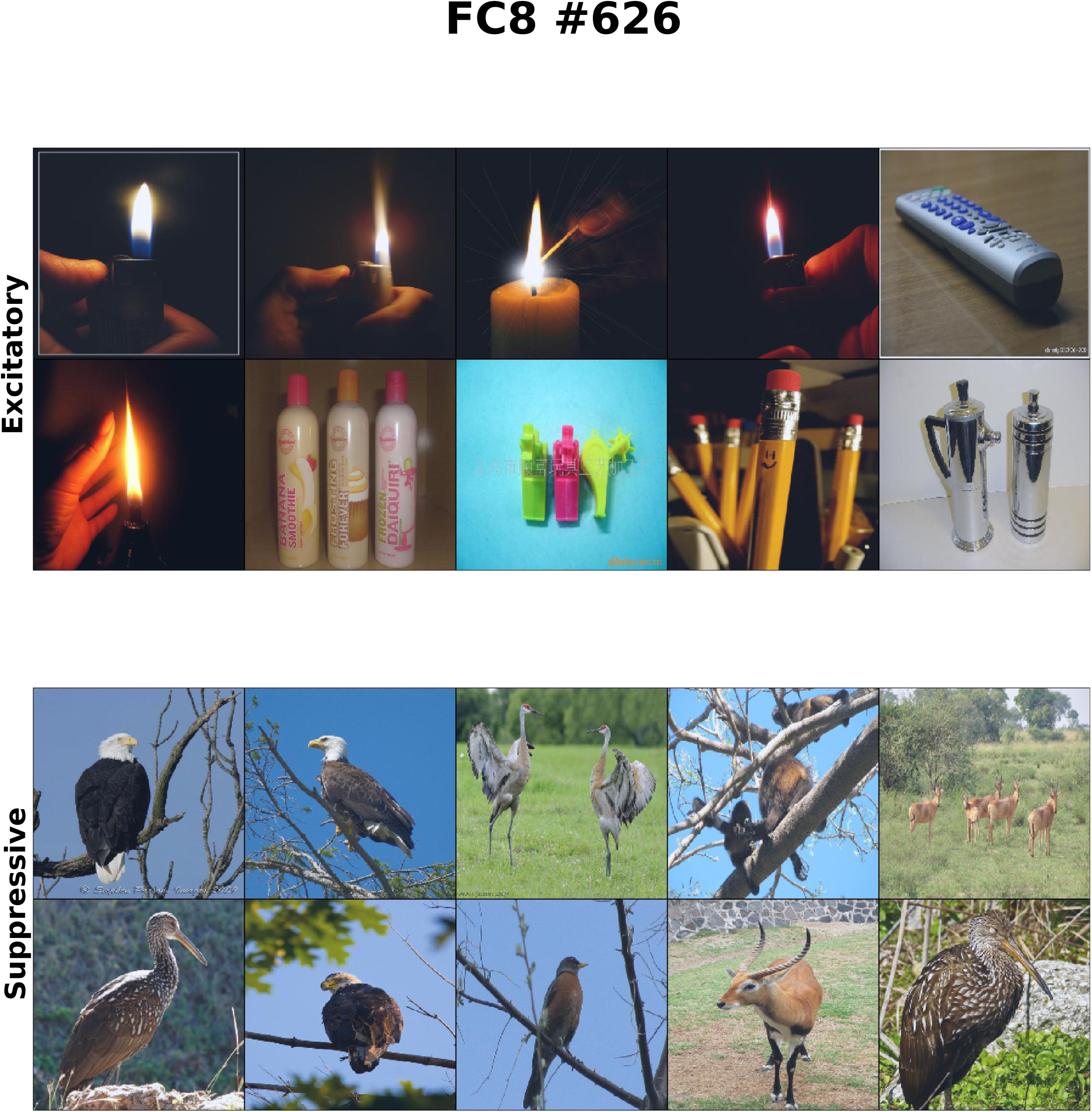
FC8 #626 - example unit selective for light flare. Category: lighter.

**Figure A29.**
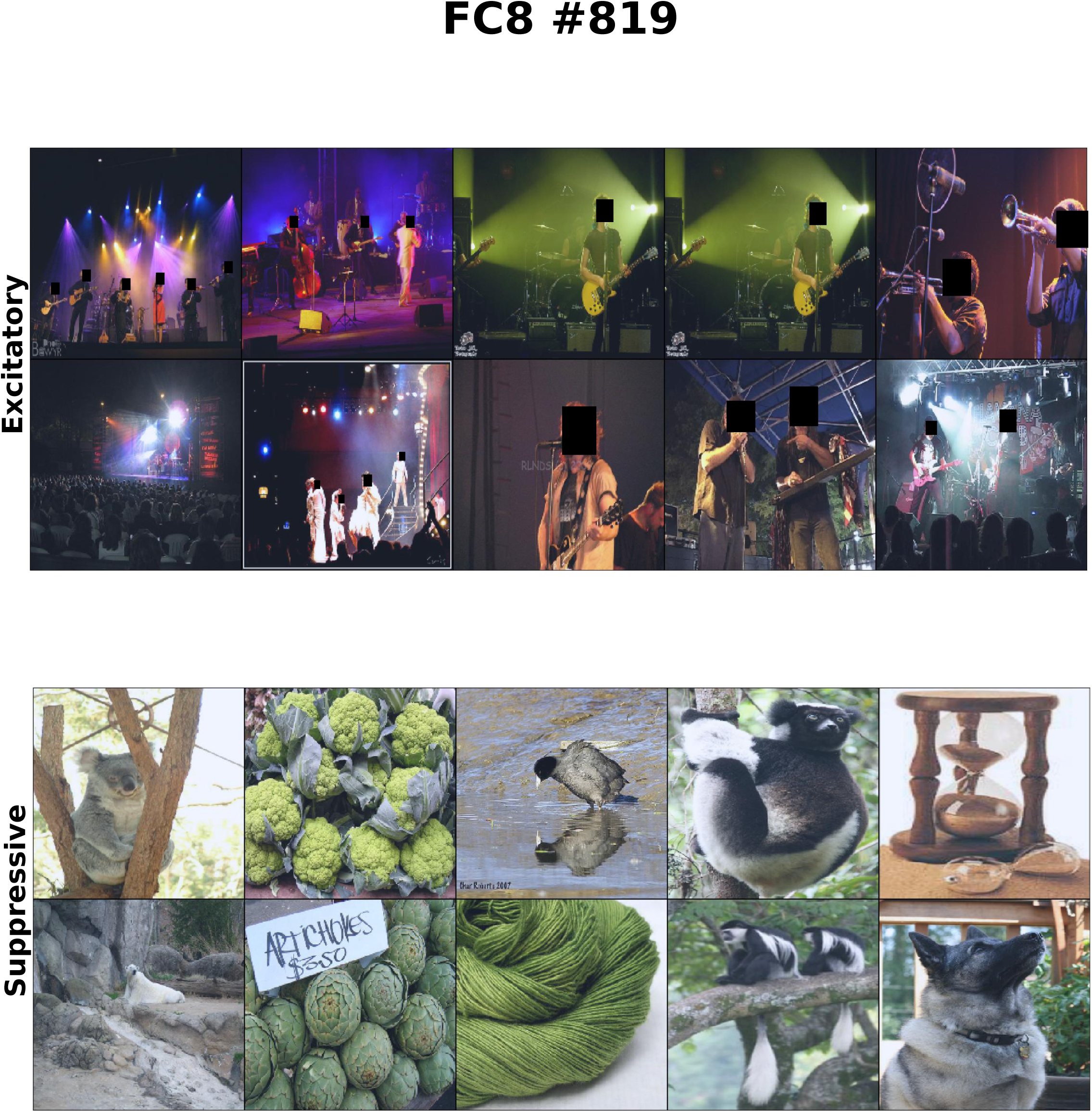
FC8 #819 - example unit selective for light flare. Category: stage.

## REFERENCES

Baker CL Jr, Mareschal I (2001) Chapter 12: Processing of second-order stimuli in the visual cortex. Progress in Brain Research

Dakin SC, Mareschal I (2000) Sensitivity to contrast modulation depends on carrier spatial frequency and orientation. Vision Research, 40: 311–329.

Dimattina D, Baker CL Jr (2019) Modeling second-order boundary perception: A machine learning approach. PLOS Computational Biology 15(3): e1006829.

El-Shamayleh Y, Movshon JA (2011) Neuronal responses to texture-defined form in macaque visual area V2. Journal of Neuroscience 31(23):8543--8555.

Elder JH (1999) Are edges incomplete? Int. J. Comput. Vision. 34, 97–122.

Held RT, Cooper EA, Banks MS (2012) Blur and disparity are complementary cues to depth. Curr Biol 22:426–431.

Jia Y, Shelhamer E, Donahue J, Karayev S, Long J, Girshick R, Guadarrama S, Darrell T (2014) Caffe: Convolutional architecture for fast feature embedding. arXiv. https://arxiv.org/abs/1408.5093.

Krizhevsky A, Sutskever I, Hinton GE (2012) ImageNet classification with deep convolutional neural networks. NIPS 25, pp 1–9.

Li S, Kwok JT, Wang Y (2001) Combination of images with diverse focuses using the spatial frequency. Information Fusion 2:169–176.

Ma K, Fu H, Liu T, Wang Z, Tao D (2018) Deep blur mapping: exploiting high-level semantics by deep neural networks. IEEE Transactions on Image Processing 27(10) 5155–5165.

Mareschal I, Baker CL Jr (1998) A cortical locus for the processing of contrast-defined contours. Nature Neuroscience 1:150–154.

Oleskiw TD, Nowack A, Pasupathy A (2018) Joint coding of shape and blur in area V4. Nature Communications 9:466.

Pasupathy A, Connor CE (2001) Shape representation in area V4: position-specific tuning for boundary conformation. J Neurophys 86:2505–2519.

Pospisil DA, Pasupathy A, Bair W (2018) ‘Artiphysiology’ reveals V4-like shape tuning in a deep network trained for image classification. eLife 2018;7:e38242.

Shapley R, Hawken MJ (2011) Color in the cortex: single- and double-opponent cells. Vision Research 51:701–717.

Zeiler MD, Fergus R (2013) Visualizing and Understanding Convolutional Networks. arXiv:13112901 [cs] Available at: http://arxiv.org/abs/1311.2901.

